# SP2G: an imaging and analysis pipeline revealing the inter and intra-patient migratory diversity of glioblastoma

**DOI:** 10.1101/2023.02.24.529679

**Authors:** Michele Crestani, Nikolaos Kakogiannos, Fabio Iannelli, Tania Dini, Claudio Maderna, Monica Giannotta, Giuliana Pelicci, Paolo Maiuri, Pascale Monzo, Nils C. Gauthier

## Abstract

Glioblastomas are heterogeneous, primary brain tumors hiding several sub-populations. Patient-derived xenografts are considered gold-standards to study glioblastoma invasion. However, they present many disadvantages, including time consumption, complex standardization, high cost. To counteract these issues and rapidly identify the most invasive sub-populations, we developed an *in vivo* mimicry platform named SP2G (SPheroid SPreading on Grids). Live imaging of tumor-derived spheroids spreading on gridded micro patterns mimicking the brain vasculature recapitulated 3D motility features observed in brain or 3D matrices. Using patient-derived samples coupled with a semi-automated macro suite, SP2G easily characterized and sorted differences in cell migration and motility modes. Moreover, SP2G exposed the hidden intra-patient heterogeneity in cell motility that correlated molecularly to specific integrins. Thus, SP2G constitute a versatile and potentially pan-cancer workflow to identify the diverse invasive tumor sub-populations in patient-derived specimens. SP2G includes an integrative tool, available as open-source Fiji macro suite, for therapeutic evaluations at single patient level.

**Teaser:** Cracking the inter and intra-patient diversity in Glioblastoma migration profiles

## Introduction

Cell migration is a complex process orchestrated by intracellular, intercellular, and environmental determinants (*1*). For the most aggressive form of glioma (i.e. grade IV glioma, or glioblastoma (GBM)), studying its motility is opportune given their extremely diffusive nature (*2*) and heterogeneity within and between patients (*3, 4*). GBM is endowed with stem cell features (*5, 6*) and only 5.0% of the patients survive after 5 years upon diagnosis (*7, 8*). Treatment options for GBM rely on surgical resection, coupled with radio- and chemotherapy, which in most cases are insufficient to cure the patient (*9, 10*). Despite these aggressive treatments, most GBMs recur a few centimeters away from the original site (*11, 12*).

GBM diffusive behavior and population heterogeneity drive GBM recurrence. The combination of these two factors represents the main hindrance to a cure. However, the interplay between heterogeneity and diffusion by migration is unclear. Moreover, the holistic dissection of GBM migration through a scalable, minimal, yet comprehensive imaging and analysis workflow remains challenging. This is because integrating the pre-existing brain structures in a reproducible and 3D-like system often precludes optical accessibility, time profitability, and analytical workflows (*13, 14*). In 1938, H.J. Scherer described (*15*) glioma cells as “replacing the nerve cells and their dendrites”, “arranging outside the Virchow-Robin spaces” (i.e., the perivascular space), and “following the surface of thick nerve bundles” (i.e., white matter tracts). These features, thereby named Scherer’s structures, intertwine in a network of topographical linear cues that foster GBM migration (*16–22*).

Considering such network complexity is crucial while designing *in vitro* systems to study GBM migration and its motility modes, as mechanical and chemical cues by the microenvironment are known to influence glioblastoma behavior and molecular signature (*23–27*). For example, as white matter tract mimicries, electrospun fibers (*28, 29*) and linear grooves (*30, 31*) provide just a single linear topographical guidance. When under confinement on printed lines mimicking blood vessels topography, glioma cells exhibited a saltatory locomotion similar to the motility modes observed *in vivo* (*32*), which were partially characterized by employing biocompatible materials that recapitulate 3D invasion (*33, 34*). Engineered vessels (*14*) and microvascular networks (*35, 36*), while potentiating GBM migration, are inadequate for broad systematic investigation due to co-culture issues that might compromise GBM stemness, complexity, and reproducibility. Cerebral organoids (*37*), organotypic brain slices (*13, 38*), and mouse xenografts (*39*), fully recapitulate the *in vivo* brain architecture and highlight the collective interconnection of invading GBM in communicating networks. However, they require laborious protocols and are inappropriate for robust analytical tool development needed to accelerate discoveries and diagnosis. Altogether, these impediments preclude a holistic dissection of GBM migration and motility modes and fail to provide a robust and broadly applicable platform to unveil how the cancer heterogeneity found in patients impacts cell invasiveness through motility along brain topographical cues.

To fulfill this most needed analytic platform, we thus developed SP2G. It is an easy and time-profitable method that integrates live cell imaging with a dedicated analysis workflow for comprehensively characterizing GBM migration and motility modes well into the intra-patient cancer heterogeneity. SP2G, using live cell imaging, combines spheroid or tumorspheres, as the best patient-derived proxy available in hospitals worldwide, with gridded micropatterns (*40*), as one of the simplest *in vitro* platforms for efficiently mimicking the brain vascular network. Biologically, spheroids maintain GBM stemness and heterogeneity (*41*) and, in SP2G, they allow to study cellular interconnections during motility, an hallmark of GBM invasion *in vivo* (*39*).

## Results

### SP2G mimics glioblastoma invasion on brain blood vessels

To validate our SP2G assay, we surveyed three-dimensional (3D) and two-dimensional (2D) techniques both with and without vessel-like topographical cues (Fig. 1 and Supplementary Video 1). For this survey, we used rat C6 glioma cells, a well-established model that migrate efficiently on host brain vasculature (*19*). We imaged <u>sp</u>heroid <u>sp</u>reading (SP2) in 5 different settings: mouse brain slices (Fig. 1A)(*42*), 3D hydrogels (Fig. 1B,C) (6mg/ml collagen I and 10mg/ml Matrigel)(*34, 43*), 2D flat substrates (laminin-coated dishes) (Fig. 1D) and laminin gridded micropatterns (SP2-G) (Fig. 1E). As observed in figure 1A-E, spheroids spread faster on 2D flat and gridded micropatterns (~8h for the complete dissolution of the spheres) compared to 3D gels (>24h) and brain slices (>48h). More importantly, time projections revealed that C6 cells aligned along the blood vessels in brain slices, while in 3D gels they invaded isotropically (Fig. 1A-C and Fig. S1D,E). Similarly, C6 cells aligned along the grid when migrating on the micropatterns, while they spread out isotropically on 2D-flat (Fig. 1D-E). We quantified spheroid spreading by measuring the areas of the spheroids at various time points (24h for 3D gels and 8h for 2D and micropatterns, Fig. 1G). Spheroid spreading in brain slices was not quantifiable due to tissue opacity and scattering of the GFP signal (Fig 1A). As observed in figure 1G, spheroid spreading was higher in matrigel than collagen (10.58 ± 0.75 and 4.76 ± 1.31, respectively; mean ± s.d.) and higher on grids than 2D flat (19.73 ± 3.81 and 14.82 ± 2.70, respectively; mean ± s.d.). Moreover, spheroid spreading was higher on laminin (LN) than fibronectin (FN), collagen (CN) or poly-L-lysine (PLL) on 2D flat (Fig. S1A,B). This confirmed laminin as the best matrix protein to study glioma motility (*28, 31, 32, 40*). The size of the linear tracks, the distance between the tracks, and the arrangement of junctional nodes were similar to the geometry of immunostained brain blood vessels. Together these results showed that our gridded micropatterns was a simple but realistic proxy to mimic brain blood vessel tracks (Fig. S1C-E).

**Fig. 1.**
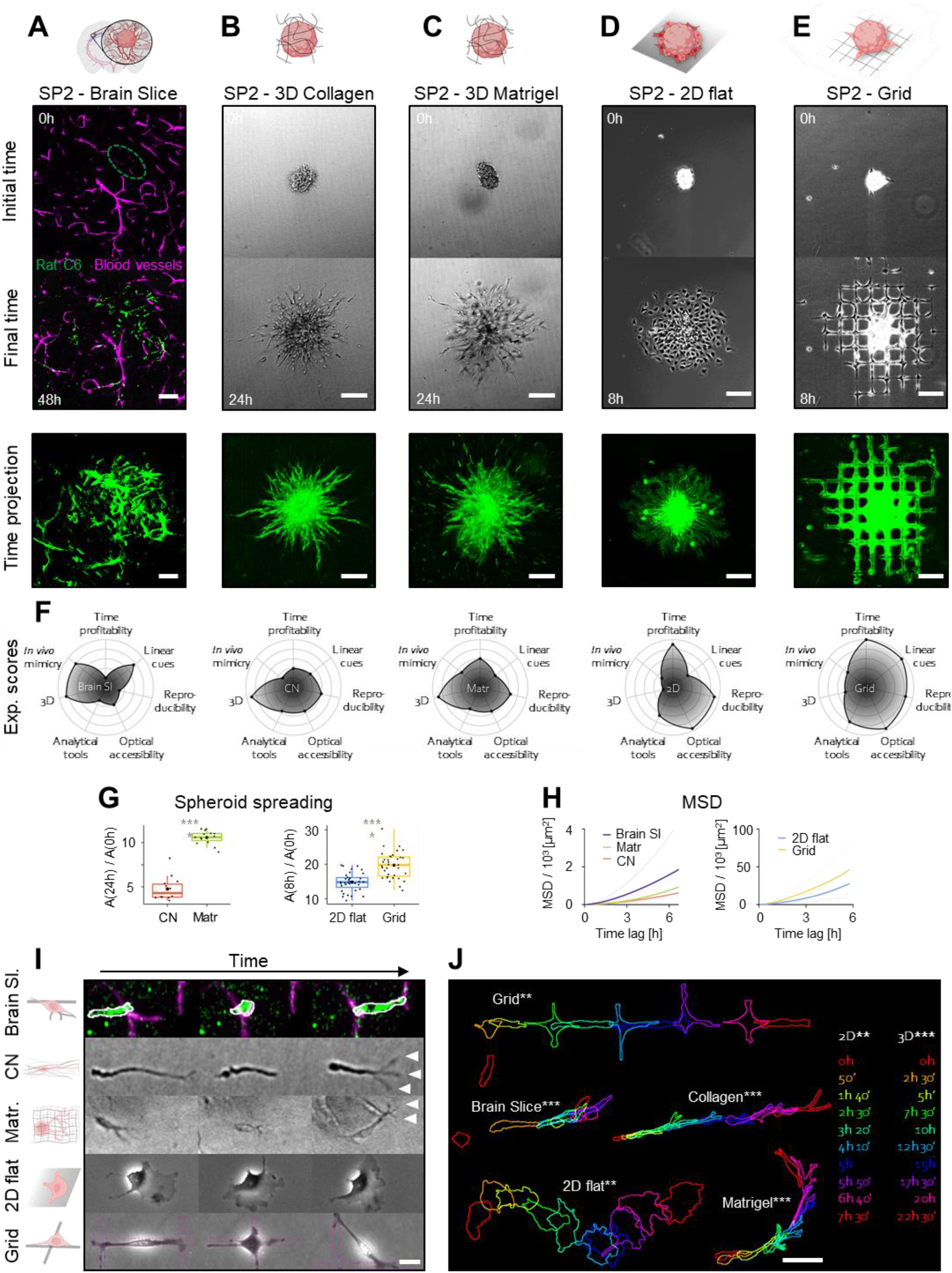
Spheroid spreading on grids (SP2G) mimics glioblastoma invasion on brain blood vessels. **(A-E)** Rat C6 glioma cells cultured as spheroid were stained (green, DiOC6 dye) and seeded on mouse brain slices (A), in collagen gel (B), in matrigel (C), on laminin-coated dishes (D), on gridded micropatterns (E) and imaged for 48h (A), 24h (B,C) or 8h (D,E). First and last images of the movies (upper panels) and time-projections (lower panels) are shown. The dashed oval corresponds to the initial area of the spheroid in the brain slice. **(F)** Radar plots summarizing experimental scores (1 to 5) of time profitability, presence of linear cues, experimental reproducibility, optical accessibility, possibility to develop analytical tools, three-dimensionality (3D) and in vivo mimicry for each setting. **(G)** Quantification of spheroid spreading in collagen gel, matrigel, 2D flat and grid (n = 14, 15, 35, 35 spheroids). Two-tailed unpaired t-test (****, p<0.0001). **(H)** Mean squared displacement (MSD) plots obtained from single cell tracks migrating in brain slice, collagen gel, matrigel, 2D flat and grid (n = 80, 95, 90, 216, 216 tracks; 5 to 7 tracks per spheroid. n = 2, 2, 2, 6, 6 independent experiments). Friedman test for Brain slice-Collagen-Matrigel (p<0.0001 in all the multiple comparisons), Mann-Whitney test for 2D-grid (p=0.0001). **(I)** Snapshots of single cells moving away from the spheroid in each setting, extracted from movie S1. **(J)** Panel summarizing cell shapes for C6 cell motility in each setting. Time is color-coded as indicated. Bars are 100 μm (A-E), 20 μm (I), and 50 μm (J)

We then compared single cell motility in the 5 settings (Fig. 1H-J and S1F-G). We tracked single cells to evaluate their migration efficiency, as Mean Squared Displacement (MSD, Fig. 1H), mean velocity (Fig. S1F-G) and cell shape (Fig. 1H,I). Cells migrated faster when the experimental setup provided vessel-like topographical cues (Brain slices and grids, Fig. S1C-E), compared to conditions devoid of linear guidance (Collagen, Matrigel and 2D flat, Figs. 1H, S1F-G). On grids and brain slices, cells showed elongated shapes and “stick-slip” motility features (*32, 44, 45*) (Fig. 1I,J and Supplementary Video 1). Conversely, in 3D hydrogels, cells protruded multiple finger-like structures, likely due to the tangled architecture of the microenvironment (Fig. 1I, arrowheads). On flat surfaces cells adopted a fan-like shape as previously described on homogeneous substrates (*32, 46*).

Moreover, we found that spheroid spreading and single cell motility were independent of the original spheroid size (Fig. S1H,I), strengthening SP2G reproducibility as spheroid sizes are usually hard to homogenize.

Finally, we analyzed the performance of each technique with radar plots using an indexing system from 1 to 5 (poorest to best performance) for 7 key parameters in GBM motility analysis: time profitability, presence of linear topographic cues allowing stick-slip motility, experimental reproducibility, optical accessibility, possibility to implement semi-automated analysis, 3D confinement and *in vivo* mimicry (Fig. 1F). We confirmed SP2G as an optimal approach to cover more requirements than other systems, with the highest scores in time and optical accessibility. SP2G preserved linear cues and junctional nodes, which are crucial in influencing motility modes, decision making for directionality and molecular signatures in different model systems (*24–27, 30, 32, 40*).

### SP2G experimental setup and image analysis workflow

To quantitatively describe cell migration and motility modes, we designed a semi-automated analysis workflow (Fig. 2A,B) composed of 7 open-source ImageJ/Fiji macros, which ultimately delivered 6 outputs: 1) migration area, 2) diffusivity, 3) boundary speed, 4) collective migration, 5) directional persistence and 6) hurdling, i.e. cell crossing the non-adhesive areas (*40*) (see Supplementary Appendix for computational details and user manual). For a better cell segmentation and a stable readout, grids and spheroids are stained with fluorescent dyes and spreading is imaged by fluorescence and phase contrast microscopy (Fig. 2A and Supplementary Video 2). We divided SP2G image analysis workflow (Fig. 2B and fig. S2A-E) in 2 main steps as follows. The first step processes the raw data semi-automatically and characterizes cell migration with the outputs # 1 (migration area, A(t)), # 2 (diffusivity, D(t)) and # 3 (boundary speed, v(t)). This is achieved by combining the binarized images of the grids and the spreading spheroids in order to construct a polygon that connects the grid nodes traveled by the spheroid invasive front at each time point. A time-lapse of a growing polygon representing the leading front of the spheroid is then automatically generated for each spheroid. An average migration area A(t) per cell line (>10 spheres per cell line) is represented by firstly aligning the baricenters of the polygons, and subsequently by extrapolating the mean xy coordinates of all the polygons at each time point (Fig. 2C, output 1). The corresponding numerical values are smoothed and differentiated in time to obtain diffusivity values D(t) (Fig. 2C, output 2). Then, boundary speed v(t) (Fig. 2C, output 3) is derived from the formula v(t) = D(t) / (2√(A(t))) (see Methods and Supplementary Appendix).

**Fig. 2.**
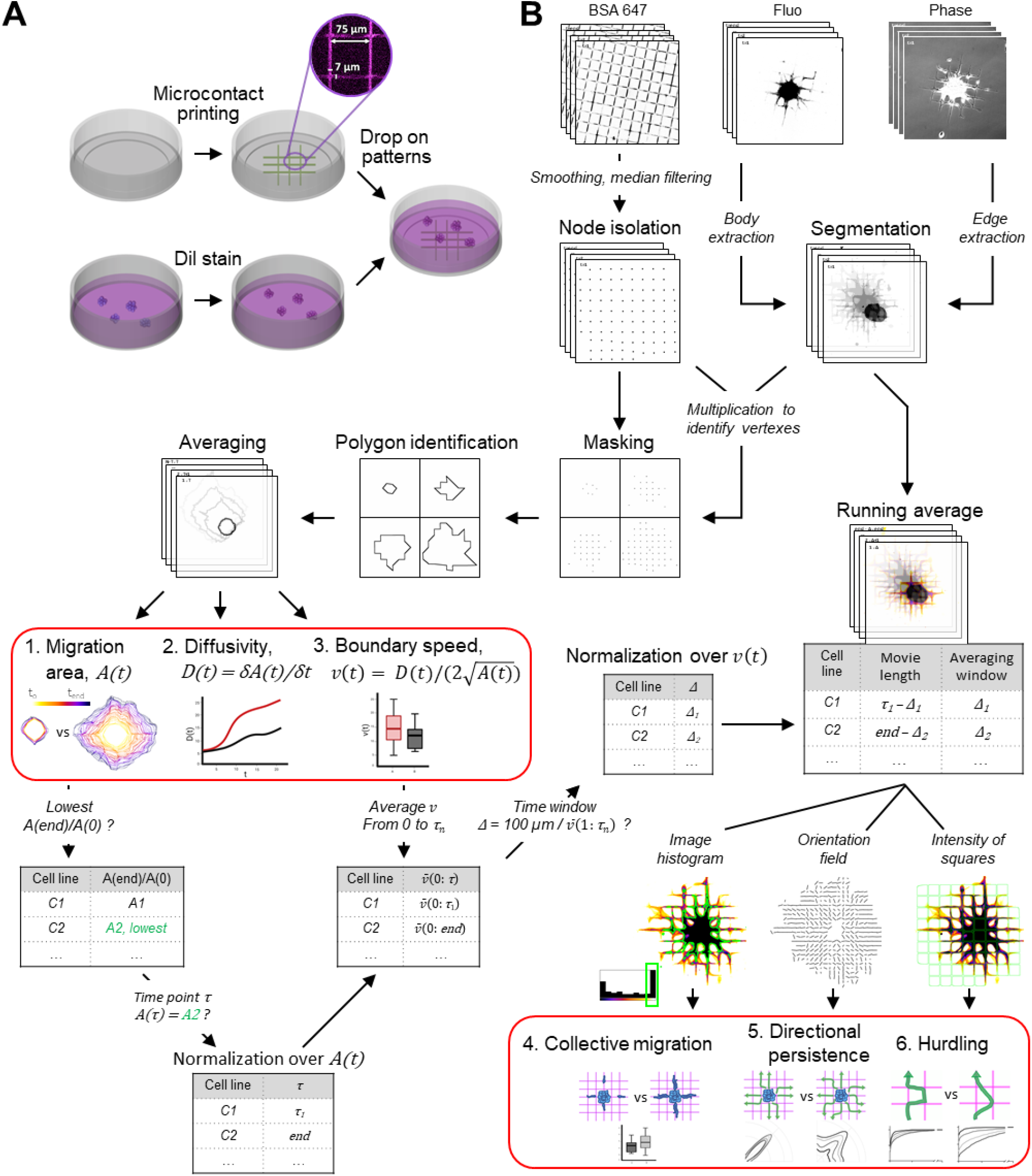
SP2G experimental setup and image analysis workflow. **(A)** Spheroids are stained (red, Dil), seeded on fluorescent gridded micro-patterns (coated with laminin mixed with BSA 647) and imaged for 8h. **(B)** Cells and grids images are segmented in binary images that are multiplied to isolate the grid nodes covered by the invasive boundary. Polygons tracking the spheroid spreading in time are then reconstructed and averaged. The time trend is projected and color-coded to visualize migration area A(t) (output #1). Diffusivity D(t) is obtained by differentiating A(t) (output #2). Spheroid boundary speed v(t) is obtained from D(t) and A(t)(output #3). Normalization of A(t) and v(t): τ, corresponding to the frame at which each cell line has migrated the same distance as the slowest cell line (C2) at the end of the acquisition (8h or more), and Δ, corresponding to the time window to complete 100 μm, are identified and all the movies are cut at τ. Running average (RA) movies are created by shifting Δ in the interval 1:τ and cell motility modes are characterized by extrapolating features from the RA movies: collective migration (output #4) is obtained by thresholding the area (outlined in green) of pixels belonging to the last bin of the histogram; directional persistence (output #5) is obtained by evaluating image orientation; hurdling (output #6) is obtained by sampling the intensities of the grid squares.

The second step of SP2G analysis provides the 3 other numerical outputs that characterize motility modes: collective migration (output #4: the higher the values, the more cells migrate as collective strands), directional persistence (output #5: the higher the values, the more cells stay on the same direction) and hurdling (output #6: the higher value the more cells are cutting angles). Only motile cells are considered for motility modes analysis and are defined by an average boundary speed higher than 100 μm / 8 h (0.21 μm/min). The characterization of motility modes is based on running average (RA) movies of the spreading spheroids. We defined the time window to complete 100 μm as Δ, which is calculated as 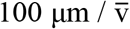, being 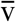 the mean v(t). When studying a stand-alone condition, RA movies are generated by shifting Δ and are long 96 (8 hours sampled every 5 min) – Δ frames. However, when comparing the motility modes of conditions with specific A(t), D(t) and v(t), SP2G requires normalization over migration area and boundary speed (see Supplementary File 1). To normalize over migration area, a time step τ has to be identified, where A(τ) / A(0) is equal to the minimum A(end) / A(0). Thus, each condition has its τ, such that A(τ) / A(0) is constant among all the conditions. Furthermore, Δ normalizes over the boundary speed and is calculated in the interval 1:τ. Each condition is endowed with its own Δ, which is larger as cells are slower. Each movie is τ – Δ frames long, covers the same area on average and each frame embeds information on the cell’s footprint in the last Δ frames.

The analysis of image features in the RA provides numerical outputs for motility modes. SP2G extrapolates collective migration by thresholding the region belonging to the last bin of the RA histogram (Fig. 2B, output 4, green area). Cells migrating collectively formed long strands protruding from the spheroid, thus giving rise to higher values when averaged in the RA. The larger this area, the higher the ratio with the total area (values are between 0 and 1). Strikingly, the numerical outputs of SP2G reflect the collective migration in simulated data of continuous, pseudo-continuous, and diffusive migration of particles at 3 speed regimes (Fig. S2F-I). SP2G computes directional persistence by evaluating the image orientation (Fig. 2B, output 5). Due to its orientation, the grid provides 2 preferential directions for the cell path: 0°, 90°. Therefore, cells capable of turning necessarily leave their footprint along 45° and 135° (the least-preferred directions), increasing their orientation values. SP2G calculates directional persistence as the ratio between orientation values in the preferential and least-preferred directions. Finally, SP2G calculates hurdling, by sampling the intensity of the grid squares (Fig. 2B, output 6; see Supplementary Excel File 2 and Supplementary Appendix). SP2G computes time trends for outputs 4-5-6, but, for the sake of simplicity, we reported their average values from the last frame of RA movies. Once calibrated and evaluated with the C6 model, we analyzed patient-derived samples.

### SP2G quantifies migratory tactics adopted by glioblastoma cells

In order to test SP2G analysis workflow, we examined the spreading of 3 patient-derived cell lines known to adopt different single-cell motility modes on grids: NNI-11 (non-motile), NNI-21 (hurdler) and NNI-24 (glider)(*40*). As observed in figure 3, SP2G confirmed their migratory behavior: within the same time window (4h) the most motile NNI-21 spread further away than the NNI-24 and NNI-11 (Fig. 3A). This behavior was reflected by a greater diffusivity (Fig. 3E, D_NNI21_(3h30’) = 1585 ± 282 μm^2^/min, D_NNI24_(3h30’) = 324 ± 204 μm^2^/min, D_NNI11_(3h30’) = 91 ± 103 μm^2^/min; mean ± s.d.) and a greater boundary speed (Fig. 3F, v_NNI21_(0:3h30’) = 1.54 ± 0.3 μm/min, v_NNI24_(0:3h30’) = 0.72 ± 0.2 μm/min, v_NNI11_(0:3h30’) = 0.19 ± 0.07 μm/min; mean ± s.d.) (Fig 3b,c,e-g). Moreover, the specific motility modes (hurdler vs glider) could be observed and quantified (Fig. 3D,H-K). Following our previous observations (*40*), NNI-21 cells migrated stochastically, with jumpy motions reflected by a low directional persistence (3.6 ± 0.7, NNI24: 8.5 ± 2.6, Fig. 3J) and high hurdling (Fig. 3K). Hurdling was visualized with cumulative distribution functions (CDF, lower slopes indicating more hurdling) and converted to numbers by dividing the average square intensity of the NNI-21 by the one of the NNI-24, returning a relative value of 2.08. NNI-24 displayed higher collective migration than NNI-21 (0.33 ± 0.07 and 0.22 ± 0.04, respectively; Fig. 3D,I and Supplementary Video 3)(*40*).

**Fig. 3.**
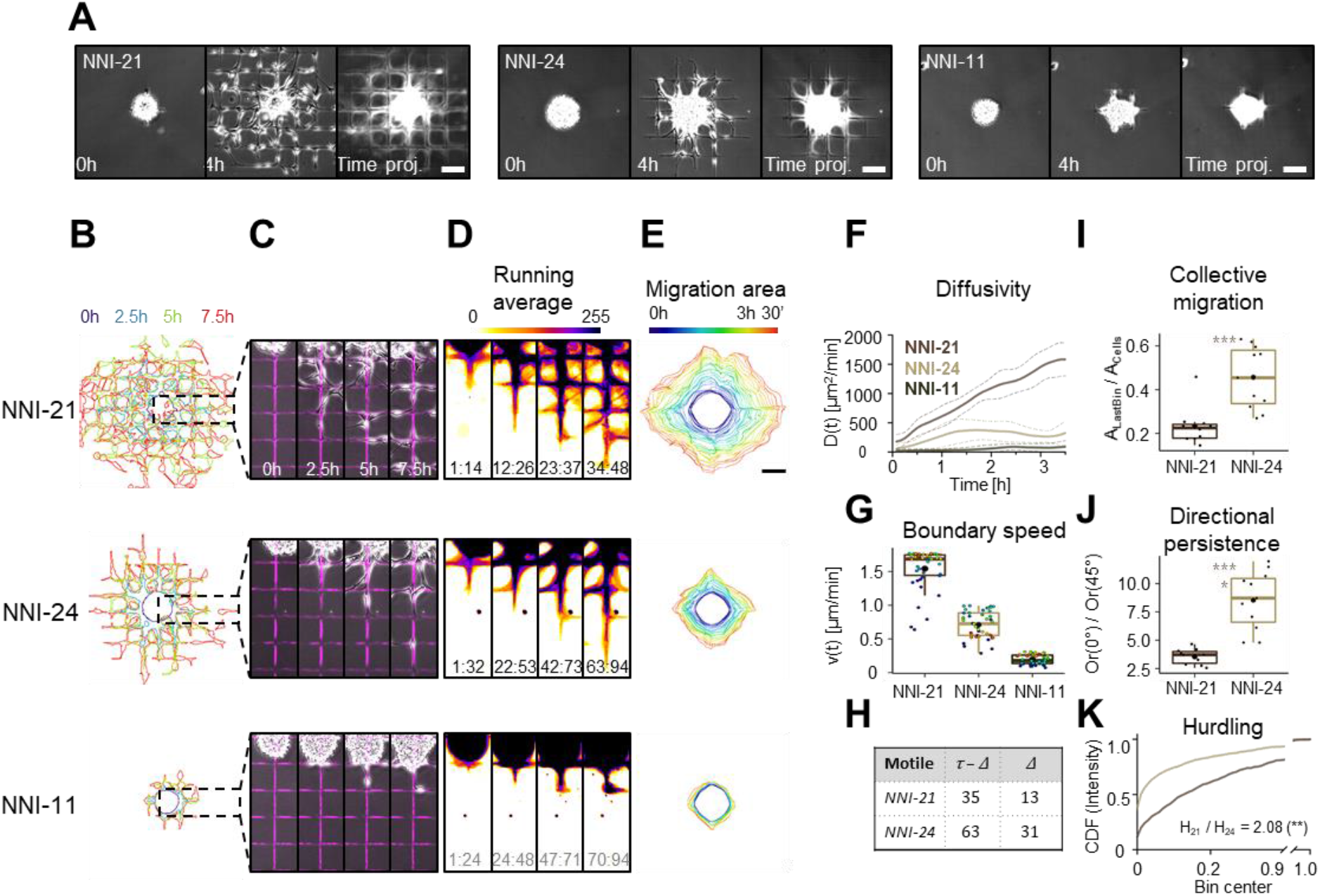
SP2G quantifies cell migratory tactics. Patient-derived glioblastoma spheroids were seeded on fluorescent gridded micropatterns, imaged for 8h, and analyzed as indicated in fig 2 (n = 10, 11, 13 spheroids, n = 2, 2, 2 independent experiments for NNI-21, NNI-24 and NNI-11 respectively). **(A)** Snapshots of the movies at 0 h, 4 h, and time projections. **(B-D)** Cellular edges (B) and corresponding overlays of the phase contrast and the fluorescent grid images (C) at 4 time points (0h, 2.5h, 5h, 7.5h) and corresponding running average (RA) (D). The time window Δ constituting the corresponding RA frame is indicated at the bottom of each panel: for the non-motile τ = 94, Δ = 24. **(E)** Average polygon visualizing migration area. **(F)** Diffusivity over 3 h 30’ (p<0.0001, Kruskal-Wallis test). Dashed lines are standard deviation. Dunn’s multiple comparison test: p<0.0001 for all. **(G)** Mean boundary speed over 3h 30’. Each dot represents a time-point and is color-coded as in (E) (p<0.0001, Kruskal-Wallis test). Dunn’s multiple comparison test: p<0.0001 for all. **(H)** Δ (number of frames needed to travel 100 μm) and τ - Δ (number of frames in the RA movie) in motile cell lines (faster than 100 μm / 8 h = 0.21 μm/min) **(I)** Collective migration for the motile cells NNI-21 and NNI-24. Each dot represents a spheroid. p=0.0004, two-tailed unpaired t-test. **(J)** Directional persistence for the motile cells NNI21 and NNI24. It is visualized as the ratio between the orientation along 0° and along 45° (Or(0°) / Or(45°), see methods). Each dot represents a spheroid. p<0.0001, two-tailed unpaired t-test. **(K)** Hurdling is visualized as the Cumulative Distribution Function (CDF) of the normalized mean intensity of the grid squares (image intensity is sampled in each square). p=0.0032 (**), Kolmogorov-Smirnov test. The ratio indicates the relationship between the average mean intensities (the sum of the mean intensity from all the squares divided by the total number of squares) of the 2 cell lines. All the bars are 100 μm. In all the boxplots, the middle horizontal line represents the median and the black dot is the mean value. Time and image intensity are color-coded as indicated.

We then evaluated SP2G sensitivity and overall performances by applying a set of cytoskeleton-perturbing drugs to NNI-21 in a dose-dependent manner. We recorded the effects for the Arp2/3 inhibitor CK666, the myosin II inhibitor blebbistatin, the microtubule poison nocodazole, and the actin poison latrunculin-A, each at 2 different concentrations (Fig. S3 and Supplementary Video 4). CK666 did not affect NNI-21 migration, whereas all the other drugs at least halved spheroid spreading (Fig. S3A-E, H). CK666 preserved cell motility modes (Fig. S3F, I, J), while blebbistatin and latrunculin-A increased collective migration and persistence, and decreased hurdling. Nocodazole kept collective migration and directional persistence but slightly decreased hurdling. Overall, these results validated SP2G as a sensitive method for motility screening and highlighted its potency as a platform for drug testing.

Cells are known to sense changes in their microenvironment (*1*), including the chemical nature of the substrate. As observed previously, GBMs have a strong tropism for laminin (*28, 31, 32, 40*). To test SP2G sensitivity towards perturbations of substrate density, we stamped the gridded micropatterns with various laminin concentrations (400, 200, 100, 50, 25, 12.5, 6.25 μg/ml) and a blank condition (no laminin) (Fig. S4). Spheroids did not adhere in the blank and at 6.25 μg/ml. Strikingly, SP2G detected 3 regimes in NNI-21 migration: 400-200 μg/ml, 100-50 μg/ml, 25-12.5 μg/ml, which were the only 3 couples non-statistically significant (Kruskal-Wallis test) when performing Dunn’s multiple comparison tests of diffusivity and boundary speed (Fig. S4E, F). SP2G measured no differences when analyzing the motility modes, except when comparing hurdling at 400 μg/ml and 12.5 μg/ml.

Altogether, SP2G emerged as sensitive in characterizing inter-patient heterogeneity in cell migration (NNI-11 vs 21 vs 24) and detected subtle motility differences under fine biochemical perturbations (NNI-21). Therefore, we hypothesized that SP2G could unveil potential intra-patient cancer heterogeneity in migration and motility modes and link this heterogeneity to specific molecular signatures.

### SP2G reveals heterogeneity in the migratory tactics adopted by glioblastoma sub-populations isolated from patient-derived tumorspheres

GBMs were characterized both as inter-(*3*) and intra-patient (*4*) heterogeneous tumors. Heterogeneous GBM displayed difference in their genomic (*47, 48*), epigenetic (*49, 50*) and transcriptomic (*51*) profiles, which were mutating under therapy (*52*) and largely maintained when cultured *in vitro* (*53*). Using SP2G, we characterized 2 patient-derived samples (the GBM7, Fig. 4 and Supplementary Video 5; and the GBM22, Fig. S5 and Supplementary Video 6).

**Fig. 4.**
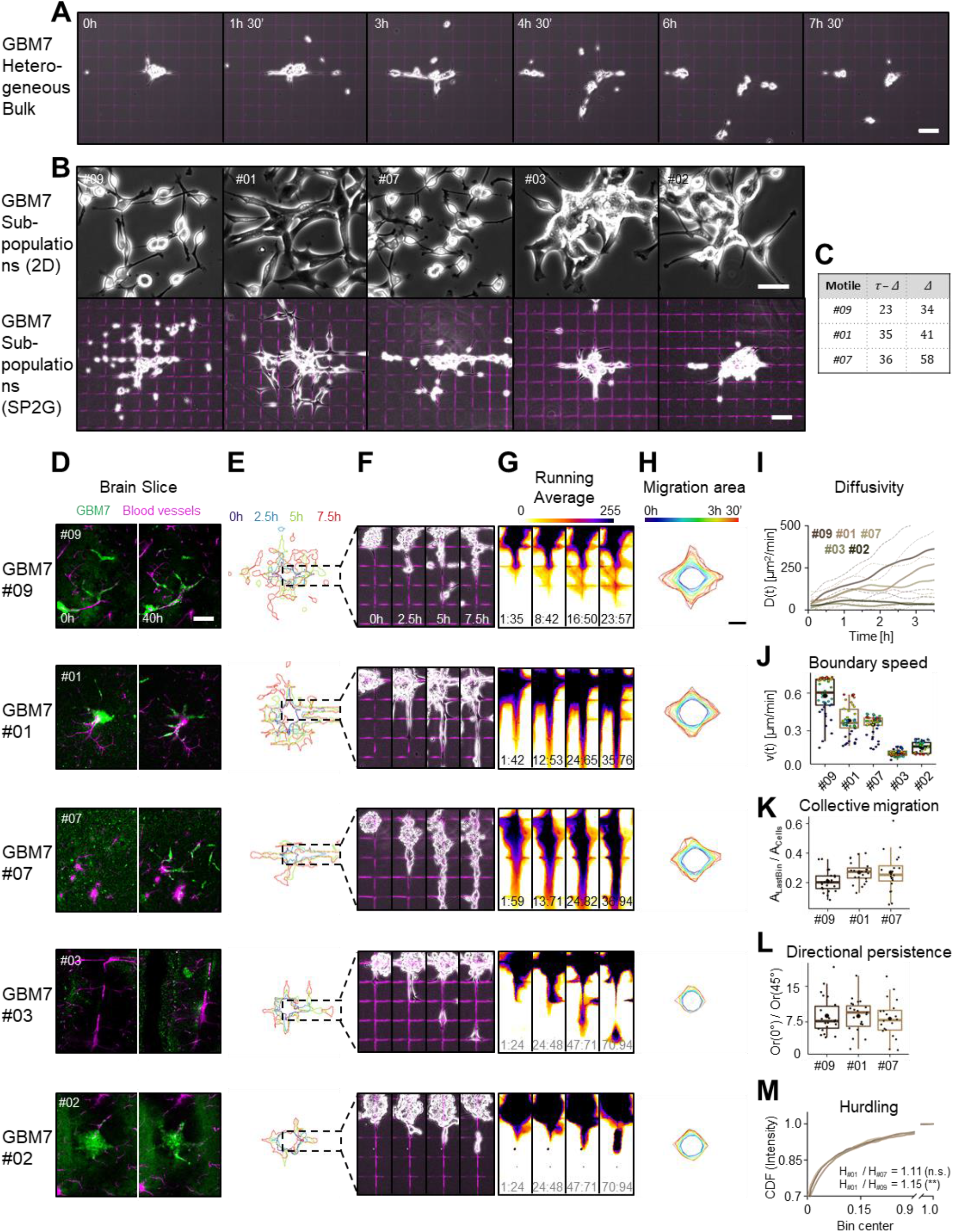
SP2G reveals migration heterogeneity in glioblastoma sub-populations isolated from patient-derived tumorspheres. Spheroids from GBM7 original cell line and isolated subpopulations (clones #01, #02, #03, #07, #09) were seeded on fluorescent gridded micropatterns, imaged for 8h, and analyzed as indicated in fig 2 **(A)** Snapshots of the movies of the original population at the indicated time-points **(B)** Phase-contrast pictures of the GBM7 sub-populations cultured on laminin (top panel, bar is 50 μm) and after SP2G at 8 hours (bottom, bar is 100 μm). **(C)** Δ and τ - Δ of the motile subpopulations. **(D)** Spheroid Spreading of GBM7 sub-populations (green, DiOC6 dye) in brain slices at 0 h and 40 h. **(E-G)** SP2G analysis of the clones #01, #02, #03, #07, #09 (n = 22, 23, 20, 22, 22 spheroids; n = 3 independent experiments): Cellular edges (E) and corresponding overlays of the phase contrast and the fluorescent grid images (F) at 4 time points and corresponding running average (RA) (G). The time window Δ constituting the corresponding RA frame is indicated at the bottom of each panel: for the non-motile τ = 94, Δ = 24. **(H)** Average polygon visualizing migration area. **(I)** Diffusivity over 3 h 30’ (p<0.0001, Kruskal-Wallis test). Dashed lines are the standard deviation. Dunn’s multiple comparison test: #09 vs #03 / #02, #01 vs #03 / #02, #07 vs #03 / #02 p<0.0001; others n.s. **(J)** Mean boundary speed over 3h 30’. Each dot represents a time-point and is color-coded as in (H) (p<0.0001, Kruskal-Wallis test). Dunn’s multiple comparison test: #09 vs #03 / #02, #01 vs #03 / #02, #07 vs #03 / #02 p<0.0001; #09 - #01 p=0.0054; #09 vs #07 p=0.0013; others n.s. **(K)** Collective migration for the motile cells #09, #01 and #07. Each dot represents a spheroid. p=0.0458, ordinary one-way ANOVA. Multiple comparisons all n.s. (#09 vs #01 p=0.0696; #09 vs #07 p=0.0903). **(L)** Directional persistence for the motile cells #09, #01 and #07. It is visualized as the ratio between the orientation along 0° and along 45° (Or(0°) / Or(45°), see methods). Each dot represents a spheroid. n.s, ordinary one-way ANOVA. Multiple comparisons all n.s. **(M)** Hurdling is visualized as the Cumulative Distribution Function (CDF) of the normalized mean intensity of the grid squares. The ratio indicates the relationship between the average mean intensities of the most hurdling (#01) against the others. The results of Kolmogorov-Smirnov tests between #01 vs #07 p=0.7992 (n.s.), #01 vs #09 p=0.0087 (**) are indicated in parenthesis. Bars are 100 μm (A, B bottom panel, D) and 50 μm (B, top panel). In all the boxplots, the middle horizontal line represents the median and the black dot is the mean value. Time and image intensity are color-coded as indicated.

Testing the GBM7 bulk, we observed a pool of motile cells tearing apart the initial spheroid into several daughter spheroids, with motile cells carrying ‘hitchhiking’ non-motile cells (Fig. 4a and Supplementary Video 5). This observation suggested that this cell line was composed of different cell populations. We isolated these sub -populations by single cell cloning and analyzed their phenotype. From the GBM7 cell line, we selected by SP2G 3 motile sub-populations (from most to least motile: #09, #01, #07) and 2 non-motile (#03, #02) (Fig. 4B,C).

The motile clones displayed different cell morphologies. Clones #09-#07 had small cell bodies and two thin long processes, whereas clone #01 had a bulkier morphology (Fig. 4B,F and Supplementary Video 5). SP2G detected lower collective migration in clone #09 compared to the other sub-populations, while clone #01 was hurdling the most. SP2G detected no significant changes in directional persistence between the 4 motile clones (Fig. 4K-M). The non-motile clones clustered in islands, with clone #03 aggregating in spheroids. In particular, clone #03 had a bulky shape and larger processes than clone #02 (Fig. 4B,F). To verify the strength of SP2G as a bona-fide alternative to brain tissues, we tested the 5 populations in the brain slice assay. The brain slice assay confirmed the results obtained by SP2G (Fig. 4D and Supplementary Video 5), with 3 motile (#09, #01, #07) and 2 non-motile (#03 and #02) clones. However, a detailed quantitative characterization of behaviors and speeds was nearly impossible with brain slice assay. Similarly, we isolated 3 motile and 2 non-motile sub-populations from the GBM22 patient-derived heterogeneous bulk (Fig. S5 and Supplementary Video 6). Clones #14-#19-#01 had small cell bodies and thin long processes, while clone #08 formed several cell-to-cell interconnections. Clones #16 and #07 grew as islands, and clone #07 clumped in spheroids (Fig. S5B,D). SP2G detected a clear trend from the most motile to the non-motile population in diffusivity and boundary speed (Fig. S5C-H). For motility modes, SP2G classified clone #08 as the most collectively migrating (Fig. S5J), clone #14 as the most hurdling (Fig. S5L), and clone #01 as the most persistent (Fig.S5K).

In conclusion, these results showed that sub-populations hidden in patient-derived samples spanned a range of migration modes comparable to those from different patients and SP2G is a valuable platform to unveil them. Next, we asked whether the transcriptional profiles of our sub-populations could account for the differences in cell motility.

### Intra-patient heterogeneity in motility modes is correlated with specific molecular signatures

We profiled transcriptional landscapes of the clones GBM7 by RNA-seq to see if their signatures correlated with their motile phenotype. Differential expression analysis followed by principal component analysis (PCA) showed that the motile (Clones #01, #07 and #09) and the non-motile (Clones #02, #03) groups clustered in 2 distinct clouds, as previously differentiated by SP2G (Fig. 5A). Gene set enrichment analysis (GSEA) of differentially expressed genes in motile versus non-motile groups showed enrichment in the ECM-receptor interaction and focal adhesion pathways (Fig. 5B) that are key pathways linked to motility. Strikingly, z-score of expression levels of integrin genes indicated that the laminin-binding integrins (particularly ITGA1, ITGA3, ITGA6) were enriched in the motile clones compared to the non-motile. Conversely, fibronectin-binding integrins were either poorly expressed or uncorrelated to cell motility (Fig. 5C). These results were confirmed by qPCR that show that the laminin-binding integrins ITGA3 and ITGA6 were highly enriched in the motile clones while the fibronectin-binding integrin ITGAV was not (Fig. 5D). These results were also confirmed by western blot (Fig. 5E). Taken together, these results validate the motile versus non-motile classification made with SP2G and provide insights on the molecular determinants that characterize GBM intra-patient heterogeneity in cell motility.

**Fig. 5.**
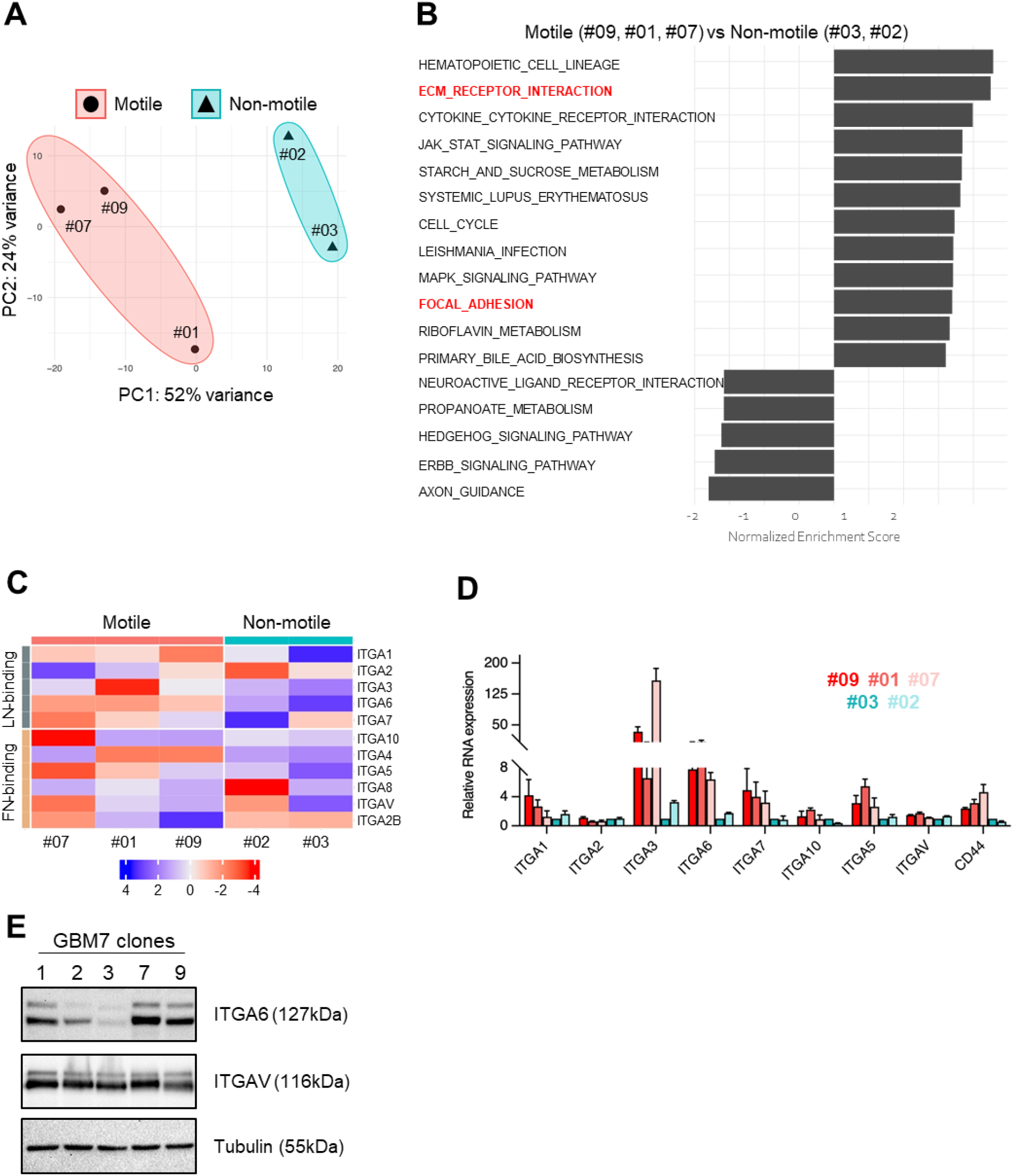
Intra-patient heterogeneity in motility modes is correlated with specific molecular signatures. **(A)** Principal component analysis showing segregation of the 5 GBM7 sub-population in motile and non-motile groups. **(B)** Gene set enrichment analysis (GSEA) of differentially expressed genes in the motile vs non-motile group. GSEA was performed using the Kyoto Encyclopedia of Genes and Genomes (KEGG) gene set in the GSEA molecular signatures database. Moderated t-statistic was used to rank the genes. Reported are Normalized Enrichment Scores (NES) of enriched pathways (with the fill color of the bar corresponding to the P-value. P-value was calculated as the number of random genes with the same or more extreme ES value divided by the total number of generated gene sets. **(C)** Heatmap representing z-score of expression levels of integrins. Laminin-binding integrins (ITGA1,2,3,6,7,10) are enriched in the motile clones. **(D)** mRNA expression levels of ITGA1, ITGA2, ITGA3, ITGA5, ITGA6, ITGA7, ITGA10, ITGAV, and CD44 determined by qRT-PCR in the 5 GBM7 sub-populations. The data are normalized over the expression in clone #03. n=3 independent experiments each. **(E)** Expression of ITGA6, ITGAV, tubulin, in total cell extracts of the 5 GBM7 sub-populations.

## Discussion

In heterogeneous tumors such as glioblastoma, some cells can be highly aggressive, migrating long distances from the tumor core by following Scherer’s structures, while other cells can be less motile and maybe more proliferative likely remaining in the tumor core. Our goal was to develop a method to rapidly define the various motility modes present in single patients and to identify which ones are the most efficient at invading mammalian brains. In previous studies, we and others, demonstrated that linear patterns were excellent proxy to mimic the brain blood vessel tracks, allowing high resolution imaging and analysis (*28–32*). In particular, gridded micropatterns allowed to easily differentiate various motility modes (*40*). Here, we improved our grid system by using spheroid spreading assays (SP2G for <u>SP</u>heroid <u>SP</u>reading on Grids). Besides allowing a faster, semi-automated analysis of the cell migration by tracking the spreading area of the spheres, this assay allows the analysis of the cell motility on a naïve, clean substrate which has never seen a glioma cell, similarly to the surface of brain blood vessels before being invaded by glioblastoma. *In vivo*, GBM exploits the surface of blood vessels as invasive highways (*2*). Being no cells in the brain further than 25 μm from a capillary (*16*), the choice of 75 μm as grid edge is a good proxy to mimic the density of the vessel network (*54*). Moreover, the topography imposed by the grid provides linear cues exploited by cells in *ex vivo* brain slice assays and, likewise, it induces the formation of invasive strands (*13, 19*) (Fig. 1A,F). Conversely, invasion is isotropic in 3D hydrogels devoid of topographical cues (*34, 43*) (Fig. 1B,C). Our gridded micropattern triggers motility modes that recapitulate the ones seen in 3D environments (*44, 55*) while keeping the advantages of simpler *in vitro* systems, such as experimental reproducibility, time profitability and amenability to high-resolution imaging techniques. Brain slice and 3D hydrogel assays recapitulate well the cues of the *in vivo* 3D environment but imaging cell moving inside these systems can be troublesome because of opacity that, in turn, prevents the development of analytical tools.

SP2G combines the experimental section with an ImageJ/Fiji toolbox tailored on it (Fig. 2). The toolbox relies on 7 macros that deliver 3 outputs for cell migration (area, diffusivity, boundary speed) and 3 outputs for the motility modes (collective migration, directional persistence, hurdling). The short duration of our experiments (8 hours) reasonably ensures the independence from cell proliferation. We utilized the patient-derived cell lines NNI-11, NNI-21 and NNI-24 as a benchmark to assess the performance of SP2G analytical toolbox (Fig. 3). In our previous work (*40*), we extensively studied the migration and the motility modes of these 3 cell lines. The NNI-11 were non-motile and highly proliferative, while the NNI-21 and NNI-24 were both diffusive, endowed with a stochastic, jumpy motion (NNI-21, named hurdlers) or following the same track with high persistence (NNI-24, named gliders). Once tested with SP2G, the differences in motility became evident. NNI-11 were non motile and NNI-21 and NNI-24 mirrored their known migration trend (90 μm/h for the NNI-21 with SP2G and 60 μm/h for single cells on grid (*40*); 40 μm/h and 30 μm/h for the NNI-24 in SP2G vs single cell on grid (*40*). SP2G identifies the hurdling of NNI-21 as opposed to the high directional persistence of NNI-24 (Fig. 3K). Furthermore, SP2G highlighted the formation of collective strands in the NNI-24, which is boosted as the spheroid acts as a reservoir for the cells to diffusively spread. Altogether these results highlighted SP2G strengths in identifying motility modes with great details and a level of refinements hard to reach with other experimental approaches.

For migration and invasion studies, spheroids are the *sine-qua-non* to dissect the motility modes behind the transition from a clustering niche towards a diffusive entity and SP2G offers a platform to tackle these questions. Spheroids also allow to study the influence of biomaterials, microenvironmental cues, and drugs sensitivity in an organoid-like configuration (*26, 33, 34, 43, 56, 57*). Our results using a panel of cytoskeleton-perturbing drugs illustrate the suitability of SP2G as a sensitive drug screening platform. We unveiled that the most invasive patient derived line, NNI-21, was dependent on myosin II, microtubule, and f-actin for migration and motility modes, while it was independent of arp2/3, similarly to previous findings (*32*). This confirmed the suitability of SP2G as a motility platform to tackle glioblastoma invasion and showed its sensitivity, as differences at low dose of drugs were visible, particularly for microtubule perturbations (Fig. S3). Moreover, SP2G gives access to area, diffusivity, boundary speed, collective migration, directional persistence and hurdling, providing an added value in the understanding of drug effects on glioblastoma behavior. By modifying the density of laminin of the grid (Fig. S4), we showed how SP2G fits in tackling micro-environmental cues like fine differences in the biochemical composition of the substrate.

SP2G underlined other key aspects of GBM migration. For example, it revealed the hitchhiking motility mode in our patient-derived samples (Fig. 4A). In our example of the GBM7 patient, the spheroid is torn apart into 3 daughter spheroids, leading us to hypothesize the presence of hidden sub-populations within a heterogeneous tumor. This behavior is kept as spheroids maintain intra-heterogeneity and stemness of the native cancer while forming a clustering niche. On one hand, the presence of glioma stem cell niches endows the cancer with plasticity that relies on stemness and heterogeneity (*3, 4, 52, 53*), but on the other hand, recent avenues propose to leverage this plasticity to induce an indolent differentiation state (i.e. lacking differentiation and tumor initiation capacity) potentially targetable with existing therapies (*58*). Indeed, heterogeneity is at the heart of potential strategies to tackle glioblastoma and, so far, it was studied only with genomic and transcriptomic tools (*47–51*), while its effect on motility was poorly understood.

Here we have reported how 11 sub-populations derived from 2 patient-derived samples behave differently in terms of migration, hurdling, persistence and collective migration (Fig. 4 and Fig. S5). The sub-populations have different cell shapes, some aggregate spontaneously in islands or spheroids when cultured in 2D, others are prone to form several cell-to-cell interconnections. Furthermore, our transcriptomic analysis (Fig. 5) showed clustering of the motile versus non-motile groups and, while looking for enrichment in gene sets key pathways, ECM-receptor interaction and focal adhesion emerged. We confirmed this result by qPCR and western blot, showing a higher expression in the laminin binding integrins, in agreement with glioma preference for laminin and with the large presence of laminin on brain blood vessels (*28, 31*). Interestingly, we found a higher protein expression in the motile GBM7 sub-populations for the integrin alpha 6 (Fig. 5E), which is known to regulate glioblastoma stem cells (*59*). At this stage, we do not know whether there is a correlation between stemness and sub-population motility, neither if the hitchhiking mode might be present *in vivo*. Tracing the history of patients by correlating with *in vivo* data is likely to help on how heterogeneity and GBM diffusivity mutually drive GBM invasion in patients.

In summary, we have presented a methodology that integrates the time-lapse imaging of spheroid spreading on grids with an ImageJ/Fiji analytical toolbox that quantitatively characterizes cell migration and motility modes. It is nicknamed SP2G, and we hope it opens up a new standard for motility screenings, potentially extendable as a pan-spheroid approach that helps answering questions on how cell migration impacts on cancer dissemination.

## Materials and Methods

### Cell culture

Rat C6 cells were cultivated in high-glucose DMEM supplemented with glutamine and 10% fetal bovine serum (FBS). To form spheroids, ~2·10^6^ C6 cells were seeded in 6-cm petri dishes previously treated for 1h with 0.2% pluronic F127 in DPBS at room temperature. After 1 day, spheroids between 75 and 150 μm in diameter were obtained. Patient-derived GBM samples from the National Neuroscience Institute (NNI-11, NNI-21 and NNI-24) were acquired with informed consent and de-identified in accordance with the SingHealth Centralised Institutional Review Board A. Patient-derived GBM samples (GBM-7, GBM-22) from the laboratory of G. Pelicci (IEO, Milan, Italy) were acquired according to protocols approved by the Institute Ethical Committee for animal use and in accordance with the Italian laws (D.L.vo 116/92 and following additions), which enforce EU 86/609 Directive (Council Directive 86/609/EEC of 24 November 1986 on the approximation of laws, regulations and administrative provisions of the Member States regarding the protection of animals used for experimental and other scientific purposes). Our patient-derived GBM cell lines were kept as previously reported(*60*). Briefly, GBM cell lines were grown in non-adherent conditions utilizing DMEM/F-12 supplemented with sodium pyruvate, non-essential amino acid, glutamine, penicillin/streptomycin, B27 supplement, bFGF (20 ng/ml), EGF (20 ng/ml), and heparin (5 mg/ml). Patient-derived GBM cell lines were passaged every 5 days. All the cell lines were maintained at 37 °C and 5% CO_2_.

### Brain slice invasion assays and staining

C57BL/6J mice were used for these studies. Both males and females (in equal proportions) within each experiment originated from different litters. All of the animal procedures were in accordance with the Institutional Animal Care and Use Committee, and in compliance with the guidelines established in the Principles of Laboratory Animal Care (directive 86/609/EEC); they were also approved by the Italian Ministry of Health.. The brain slice assay was performed as reported in Er et al.(*61*) and Polleux and Ghosh(*42*). Prior to sacrifice, mice were anesthetized, their chest was cut and intra-cardiac injection was performed with 5 ml solution of Dil stain (ThermoFisher D282) to label the luminal side of blood vessels. The Dil stain was diluted 0.5 mg/ml in 100% ethanol and this solution was further diluted 1:10 in a 30% w/v solution of sucrose-DPBS. Brains were then isolated in ice-cold CaCl2^+^/MgCl2^+^ 1X HBSS (Euroclone ECB4006) supplemented with 2.5mM HEPES (complete HBSS). Brains were sectioned in 150 or 100 μm thick slices using a Leica VT1200S vibratome and placed in a glass bottom 24-well, which was previously coated at 37 °C overnight using a solution of 12.5 mg/ml laminin and 12.5 mg/ml Poly-L-lysine (1 slice/well). Slices were left 3h at 37 °C and 5% CO_2_ to consolidate on the substrate. Subsequently, glioma spheroids were gently added and the co-culture was kept 4h at 37 °C and 5% CO_2_ prior to imaging. Movies were recorded overweekend on a confocal SP5 microscope equipped with temperature, humidity, and CO_2_ control utilizing a 20X air objective (1 frame/15 min for rat C6, 1 frame/30 min for GBM-7 sub-populations). All the brain slice live experiments were performed with brain slice culture medium (68% L-glutamine supplemented DMEM, 26% complete HBSS, 5% FBS, 1% Penicillin-Streptomycin). For immunofluorescence staining of C6 and blood vessels, the co-culture was then fixed with 4% PFA for 20 min and incubated for 1h at room temperature with a blocking solution made of 5% BSA, 5% Normal-Donkey-Serum (NDS) and 0.3% Triton-X 100 in DPBS. The co-culture was incubated overnight at 4 °C with 10 mg/ml of Tomato Lectin (Vector Laboratories, #DL-1178) in blocking solution. DAPI was put afterwards. Images were acquired with a Leica SP8 microscope utilizing a 63X oil objective (1 μm Z step).

### Collagen and matrigel invasion assays

A previous protocol was adapted for collagen and matrigel assays(*62*). Briefly, 20 ml of polydimethylsiloxane (PDMS; Sylgard 184 Dow Corning) were casted at 1:10 ratio by mixing curing agent and silicone elastomer base, respectively, in a 10-cm plate. 6-mm PDMS wells were obtained by punching holes with a biopsy puncher in 18×18 mm PDMS squares. The PDMS wells were bound on a 24 mm coverslip via plasma treatment (90 s) followed by 5 min at 80 °C. Each 6-mm well was then treated with 1 mg/ml poly-d-lysine at 37 °C for 3h, rinsed in milliQ water, and cured overnight at 80 °C. Meanwhile, rat C6 spheroids were incubated in medium with 5 μM Dil stain for 3 h, then collected, centrifuged at 500 rpm for 2 minutes and re-suspended in 1 ml of medium. 10 μl of spheroid suspension was then mixed with 80 μl of 6 mg/ml collagen solution (Collagen I from rat tail, Corning #354249 diluted in cell culture medium, 10% v/v 1.2% NaHCO_3_, 5% 1M HEPES, 1.5% 1M NaOH) or 10 mg/ml Matrigel (reconstituted basement membrane, rBM, Trevigen # 3445-005-01). The spheroids embedded in unpolymerized solutions were placed in the 6-mm PDMS wells and left at 37 °C for 1h to polymerize. With this method between 5 and 15 spheroids per well were obtained. Afterwards, medium was added and movies of invading spheroids were acquired on an IX83 inverted microscope (Olympus) equipped with a Confocal Spinning Disk unit, temperature, humidity, and CO_2_ control. A 10X objective was utilized, along with an IXON 897 Ultra camera (Andor) and OLYMPUS cellSens Dimension software. Movies were obtained for > 24 hours (1 frame / 15 min) with 7.5mm Z step for RFP and DIC channels.

### Quantification of SP2 with area ratio and MSD

For SP2 in collagen and matrigel assays, SP2 was obtained as the ratio between the area occupied by spheroids at 24 and 0 hours. The area was calculated as the maximum intensity projection firstly in Z, then in time, of the fluorescent channel. For SP2 in 2D flat and grid, the ratio between the areas at 8 and 0 hours was calculated, and the areas were obtained utilizing the maximum intensity projection in time. For MSD calculation we utilized a published protocol(*63*). To get XY coordinates overtime, manual tracking was performed with the dedicated plugin in Fiji. For brain slice, collagen and Matrigel cells were followed for 14 h respectively considering as initial time point 24 h, 10 h and 10 h to fully visualize single cells.

### Microcontact printing

Microcontact printing was performed as we described (*60*). Briefly, we casted 1:10 PDMS from a dedicated silicon mold, cut it into 1×1 or 1×2 cm^2^ stamps, and coated with 50 μg/ml laminin (ThermoFisherScientific, 23017015) in DPBS for 20 min. Each stamp was then air-blow dried, leant on a 35-mm dish, then gently removed. In case of a plastic dish, the surface was then passivated with 0.2% pluronic F127 in DPBS at room temperature for 1 h, whereas for glass poly-l-lysine-grafted polyethylene glycol (0.1 mg/ml, pLL-PEG, SuSoS) was utilized. Dishes were then rinsed 4 times with DPBS and kept in medium until spheroids were seeded. For printing laminin concentrations from 400 to 6.25 μg/ml, a sequence of 6 serial dilutions (1:1 in DPBS) was carried out.

### SP2G experimental and image analysis workflow

In SP2G experimental section, the laminin solution for the gridded micropattern was mixed with 7 μg/ml BSA-conjugated-647 in order to visualize it. Similarly, 1-day old or 5-days old spheroids (for rat C6 or human patient-derived glioma, respectively) were incubated in medium with 5 μM Dil stain for 3 hours in 6-well plates previously passivated with 0.2% pluronic F127. Spheroids were then deposited on the grid and the samples were placed under the microscope and left 5 to 15 min to equilibrate. Afterwards, 8-hours time-lapse movies were recorded using a 10X objective mounted on a Leica AM TIRF MC system or onto an Olympus ScanR inverted microscope (1 frame / 5 min). 3 channels per time point (phase contrast, Dil stain fluorescence for the cells, BSA-647 fluorescence for the grid) were acquired in live cell imaging for the experiments in Fig. 3 and Extended Data Fig. 4. 2 channels per time point were acquired in the other experiments, since the grid fluorescence was recorded for just 1 frame before and 1 frame after the time-lapse movie. For the Experiments in Extended Data Fig. 4 cells were not fluorescently labeled and Dil stain channel was not acquired, the drugs were injected 25 min after imaging onset. For the characterization of cell migration, we utilized the SP2G analytical toolbox that measured the polygonal area A(t). Briefly, as SP2G formed the polygon to track the invasive boundary, the code initially multiplied the binarized grid nodes with the binarized spreading spheroid in order to obtain only the node traveled by at least 1 cell. Then, SP2G iteratively checked whether a node has blinked for at least 3 consecutive time frames. If this condition is met, the node is added to the polygon. Therefore, SP2G always stopped tracking 2 frames earlier than the total duration of any movie. Afterwards, from A(t) we derived the diffusivity D(t) = dA(t) / dt, where dt was the time frame in the time-lapse movies (5 min) and dA(t) was the difference between 2 polygonal areas at subsequent time steps. For the calculation of single cell velocity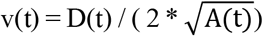, being now 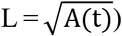 the edge of the square having an area equivalent to the polygon, the following 2-equation system was solved:

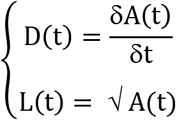

That inferred

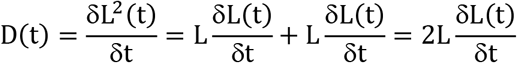

The following was obtained

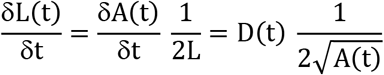

that corresponded to the value of the boundary speed. In this way a length gradient was inferred from an area gradient.

For the characterization of the motility modes we extrapolated all the parameters from RA movies and averaged data from several spheroids. Collective migration values were obtained by thresholding each frame of the RA within the last histogram bin, that necessarily spans up to 255 (the maximum value of an 8-bit image). The ratio

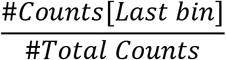

Returns the collective migration. The rationale behind this assumption was that collective strands generated high intensity values when averaged, since many cells travelled the same path. Therefore, in the RA movie there were zones of high intensity. Vice versa, single entities generated low intensities when averaged, since no cells other than the single one contributed to the final average value. Background pixels were set to NaN (see Supplementary Appendix).

Directional persistence is calculated through the function “OrientationJ distribution” of the OrientationJ plugin(*64*), which returns the orientation field (OF): it consists in 180 values (1 per direction, sampled every 1°). Reasonably, we assumed that the spheroid spreading is isotropic, and therefore SP2G averages the values 0-90, 1-91, etc. and gets 90 values. The following ratio

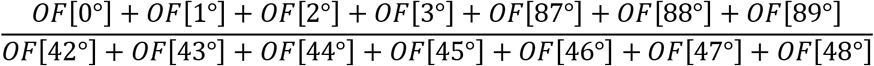

Returns the directional persistence. It is the ratio between the direction of least resistance to cell migration (i.e. the ones provided by the grid segments) and the direction of most resistance (the one a cell has necessarily to face when undergoing a directional change).

### Simulation of particle diffusion

Simulated data were generated with a custom-written code in imageJ/Fiji. Briefly, the function

> Speed_particle = speed*(1+random(“gaussian”));

Was applied at each time step to generate motion. “speed” was equal to 2, 2.5 or 3 and “random(“gaussian”)” returned a Gaussian distributed pseudorandom number with mean 0 and standard deviation 1. Continuity was set by imposing a 100% overlap probability to moving particles, pseudo-continuity a 99% probability, pure diffusivity with no constraints.

### RNA sequencing

For the RNA analysis, all the cell lines were plated on laminin at 10 μg/ml and lysed at 70-90% confluency. RNA was extracted with the RNAeasy Mini Kit (Qiagen) as per manufacturer specifications. RNA concentration was measured using Qubit 4.0 and its integrity evaluated with an Agilent Bioanalyzer 2100 utilizing Nano RNA kit (RIN > 8). An indexed-fragment library per sample was arranged from 500ng totalRNA using Illumina Stranded mRNA Prep ligation kit (Illumina) as per manufacturer’s instructions. Libraries were checked for proper size using Agilent Bioanalyzer 2100 High Sensitivity DNA kit, then normalized and equimolarly pooled to perform a multiplexed sequencing run and quantified with Qubit HS DNA kit. As a positive control, 5% of Illumina pre-synthesized PhiX library was incorporated in the sequencing mix. Sequencing was carried out in Paired End mode (2×75nt) with an Illumina NextSeq550Dx, generating on average 60 million PE reads per library. Reads were aligned with the STAR aligner (v 2.6.1d) to the GRCh38/hg38 assembly human reference genome.

Differential gene expression analysis was accomplished with the Bioconductor package DESeq2 (v 1.30.0) that assessed variance-mean dependence in count data from sequencing data. Besides, the differential expression was evaluated by employing a negative binomial distribution-based model with DESeq2. The Bioconductor package fgsea (v 1.16.0) and GSEA software (including Reactome, KEGG, oncogenic signature and ontology gene sets available from the GSEA Molecular Signatures Database, https://www.gsea-msigdb.org/gsea/msigdb/genesets.jsp?collections) were utilized for preranked gene set enrichment analysis (GSEA) to assess pathway enrichment in transcriptional data.

### RNA extraction and qPCR analysis

For the qPCR analysis, all the cell lines were plated on laminin at 10 μg/ml and lysed at 70-90% confluency. RNA was extracted with the RNAeasy Mini Kit (Qiagen) as per manufacturer specifications. Real-time PCR performed with a 7500 Real-Time PCR System (Thermo Fisher). The following parameters were utilized: pre-PCR step of 20 s at 95 °C, followed by 40 cycles of 1 s at 95 °C and 20 s at 60 °C. Samples were amplified with primers and probes for each target, and for all the targets, one NTC sample was run. Raw data (Ct) were analyzed with Excel using the DDCT method to calculate the relative fold gene expression. DCT were calculated using GAPDH as housekeeping gene and averaged (3 independent experiments). For the mRNA expression of selected integrins data were normalized against the expression of the GBM7 sub-population #03.

### Analysis of RNA sequencing data

Reads were aligned to the GRCh38/hg38 assembly human reference genome using the STAR aligner (v 2.6.1d)(*65*) and reads were quantified using Salmon (v1.4.0)(*66*). Differential gene expression analysis was performed using the Bioconductor package DESeq2 (v 1.30.0)(*67*) that estimates variance-mean dependence in count data from high-throughput sequencing data and tests for differential expression exploiting a negative binomial distribution-based model. Pre-ranked GSEA for evaluating pathway enrichment in transcriptional data was carried out using the Bioconductor package fgsea(*68*), taking advantage of the Kyoto Encyclopedia of Genes and Genomes (KEGG) gene set available from the GSEA Molecular Signatures Database (https://www.gsea-msigdb.org/gsea/msigdb/genesets.jsp?collection).

### Protein extraction and western blots

Total cell extracts were obtained utilizing RIPA buffer (100 mM NaCl; 1 mM EGTA; 50 mM Tris pH7.4; 1% TX100) complemented with a cocktail of protease inhibitors (Roche). The Pierce BCA protein assay kit #23225 (ThermoScientific) was utilized to quantify proteins, which were then denatured with SDS and resolved by SDS-PAGE using typically 8% acrylamide gels. Transfers were done on Polyvinylidene Fluoride (PVDF) membranes in methanol-containing transfer buffer. Blocking was done with milk diluted to 5% in PBS-0.1% tween for 1h at room temperature and antibodies were blotted overnight at 4 °C. HRP-secondary antibodies were incubated for 1-2 hours at room temperature in 5% milk and ECL were performed using the Amersham ECL Western Blotting Detection Reagents (Cat.no. RPN2106 from GE Healthcare). Detection was done using CL-Xposure films (Cat.no. 34089 from Thermoscientific). The following antibodies were used: anti-integrin alpha V (Abcam ab179475, diluted 1:1000), anti-integrin alpha 6 (Novus NBP1-85747, diluted 1:500).

### Statistical analysis

All the statistical analysis was performed with Prism 9 (GraphPad). P-values were calculated as indicated in figure legends, as well as number of samples and independent experiments. The plots were generated with Prism 9 and ggplot2. Google sheets was utilized for the radar plots in Fig. 1. Kolmogorov-Smirnov tests were performed on Matlab utilizing the function kstest2(groupA,groupB,’Alpha’,0.05). All the SP2G macros and the supplementary appendix containing detailed instructions on installation and run are freely available on the repository figshare at the following link: https://figshare.com/projects/SP2G/148246.

## Acknowledgments

We are grateful to the IFOM imaging facility personnel, in particular D. Parazzoli for technical support, the IFOM cell culture facility personnel, the MBI microfabrication facility personnel, in particular Sree Vaishnavi Sundararajan. We thank Marco Foiani and Giorgio Scita (IFOM), Virgile Viasnoff (MBI), Marc-Antoine Fardin (Institut Jacques Monod), Nir Gov (Weizmann Institute of Science, Israel), Scita’s and Maiuri’s groups for helpful discussions and critical comments on the manuscript.

## Funding

This work was supported by:

IFOM (starting package to NCG),

Italian Association for Cancer Research (AIRC) Investigator Grant (IG) 20716 to NCG

Italian Association for Cancer Research (AIRC) Three-year fellowship”MilanoMarathon - oggicorroperAIRC” - Rif. 22461 to MC,

MC is a PhD student within the European School of Molecular Medicine (SEMM).

## Author contributions

M.C., N.C.G. and P.M. conceptualized the research project.

M.C. and P.M. developed the computer codes and algorithms.

M.C., P.M., and N.C.G. analyzed the data.

M.C., N.K. T.D., C.M., M.G., and P.M. performed the experiments.

F.I. analyzed the RNAseq data

N.C.G. and G.P. provided the resources.

M.C. wrote the original draft.

M.C., P.M., and N.C.G. reviewed and edited the manuscript.

N.C.G. supervised the research activity.

## Competing interests

Authors declare that they have no competing interests.

## Data and materials availability

All data are available in the main text or the supplementary materials.

## Notes

https://figshare.com/projects/SP2G/148246

### Competing Interest Statement

The authors have declared no competing interest.

## References

1. P. Friedl, S. Alexander, Cancer invasion and the microenvironment: plasticity and reciprocity. Cell 147, 992–1009 (2011).

2. V. A. Cuddapah, S. Robel, S. Watkins, H. Sontheimer, A neurocentric perspective on glioma invasion. Nat Rev Neurosci 15, 455–465 (2014).

3. C. W. Brennan, R. G. Verhaak, A. McKenna, B. Campos, H. Noushmehr, S. R. Salama, S. Zheng, D. Chakravarty, J. Z. Sanborn, S. H. Berman, The somatic genomic landscape of glioblastoma. Cell 155, 462–477 (2013).

4. A. P. Patel, I. Tirosh, J. J. Trombetta, A. K. Shalek, S. M. Gillespie, H. Wakimoto, D. P. Cahill, B. V. Nahed, W. T. Curry, R. L. Martuza, Single-cell RNA-seq highlights intratumoral heterogeneity in primary glioblastoma. Science 344, 1396–1401 (2014).

5. R. Galli, E. Binda, U. Orfanelli, B. Cipelletti, A. Gritti, S. De Vitis, R. Fiocco, C. Foroni, F. Dimeco, A. Vescovi, Isolation and characterization of tumorigenic, stem-like neural precursors from human glioblastoma. Cancer Res 64, 7011–7021 (2004).

6. S. K. Singh, I. D. Clarke, M. Terasaki, V. E. Bonn, C. Hawkins, J. Squire, P. B. Dirks, Identification of a cancer stem cell in human brain tumors. Cancer Res 63, 5821–5828 (2003).

7. D. N. Louis, A. Perry, G. Reifenberger, A. von Deimling, D. Figarella-Branger, W. K. Cavenee, H. Ohgaki, O. D. Wiestler, P. Kleihues, D. W. Ellison, The 2016 World Health Organization Classification of Tumors of the Central Nervous System: a summary. Acta Neuropathol 131, 803–820 (2016).

8. Q. T. Ostrom, H. Gittleman, P. Liao, C. Rouse, Y. Chen, J. Dowling, Y. Wolinsky, C. Kruchko, J. Barnholtz-Sloan, CBTRUS statistical report: primary brain and central nervous system tumors diagnosed in the United States in 2007-2011. Neuro Oncol 16 Suppl 4, iv1–63 (2014).

9. A. Giese, R. Bjerkvig, M. E. Berens, M. Westphal, Cost of migration: invasion of malignant gliomas and implications for treatment. J Clin Oncol 21, 1624–1636 (2003).

10. L. C. Hou, A. Veeravagu, A. R. Hsu, V. C. Tse, Recurrent glioblastoma multiforme: a review of natural history and management options. Neurosurg Focus 20, E5 (2006).

11. M. E. Davis, Glioblastoma: Overview of Disease and Treatment. Clin J Oncol Nurs 20, S2–8 (2016).

12. M. Lacroix, D. Abi-Said, D. R. Fourney, Z. L. Gokaslan, W. Shi, F. DeMonte, F. F. Lang, I. E. McCutcheon, S. J. Hassenbusch, E. Holland, K. Hess, C. Michael, D. Miller, R. Sawaya, A multivariate analysis of 416 patients with glioblastoma multiforme: prognosis, extent of resection, and survival. J Neurosurg 95, 190–198 (2001).

13. T. Ohnishi, H. Matsumura, S. Izumoto, S. Hiraga, T. Hayakawa, A novel model of glioma cell invasion using organotypic brain slice culture. Cancer Res 58, 2935–2940 (1998).

14. C. Wang, J. Li, S. Sinha, A. Peterson, G. A. Grant, F. Yang, Mimicking brain tumor-vasculature microanatomical architecture via co-culture of brain tumor and endothelial cells in 3D hydrogels. Biomaterials 202, 35–44 (2019).

15. H. Scherer, Structural development in gliomas. The American Journal of Cancer 34, 333–351 (1938).

16. N. J. Abbott, A. A. Patabendige, D. E. Dolman, S. R. Yusof, D. J. Begley, Structure and function of the blood-brain barrier. Neurobiol Dis 37, 13–25 (2010).

17. G. P. Cribaro, E. Saavedra-López, L. Romarate, I. Mitxitorena, L. R. Díaz, P. V. Casanova, M. Roig-Martínez, J. M. Gallego, A. Perez-Vallés, C. Barcia, Three-dimensional vascular microenvironment landscape in human glioblastoma. Acta neuropathologica communications 9, 1–20 (2021).

18. A. P. Di Giovanna, A. Tibo, L. Silvestri, M. C. Mullenbroich, I. Costantini, A. L. Allegra Mascaro, L. Sacconi, P. Frasconi, F. S. Pavone, Whole-Brain Vasculature Reconstruction at the Single Capillary Level. Sci Rep 8, 12573 (2018).

19. A. Farin, S. O. Suzuki, M. Weiker, J. E. Goldman, J. N. Bruce, P. Canoll, Transplanted glioma cells migrate and proliferate on host brain vasculature: a dynamic analysis. Glia 53, 799–808 (2006).

20. J. Holash, P. Maisonpierre, D. Compton, P. Boland, C. Alexander, D. Zagzag, G. Yancopoulos, S. Wiegand, Vessel cooption, regression, and growth in tumors mediated by angiopoietins and VEGF. Science 284, 1994–1998 (1999).

21. F. Winkler, Y. Kienast, M. Fuhrmann, L. Von Baumgarten, S. Burgold, G. Mitteregger, H. Kretzschmar, J. Herms, Imaging glioma cell invasion in vivo reveals mechanisms of dissemination and peritumoral angiogenesis. Glia 57, 1306–1315 (2009).

22. S. Watkins, S. Robel, I. F. Kimbrough, S. M. Robert, G. Ellis-Davies, H. Sontheimer, Disruption of astrocyte-vascular coupling and the blood-brain barrier by invading glioma cells. Nat Commun 5, 4196 (2014).

23. J. Cha, P. Kim, Biomimetic strategies for the glioblastoma microenvironment. Frontiers in Materials 4, 45 (2017).

24. I. Manini, F. Caponnetto, A. Bartolini, T. Ius, L. Mariuzzi, C. Di Loreto, A. P. Beltrami, D. Cesselli, Role of Microenvironment in Glioma Invasion: What We Learned from In Vitro Models. Int J MolSci 19, (2018).

25. M. C. de Gooijer, M. Guillen Navarro, R. Bernards, T. Wurdinger, O. van Tellingen, An Experimenter’s Guide to Glioblastoma Invasion Pathways. Trends Mol Med 24, 763–780 (2018).

26. K. J. Wolf, J. Chen, J. D. Coombes, M. K. Aghi, S. Kumar, Dissecting and rebuilding the glioblastoma microenvironment with engineered materials. Nature Reviews Materials 4, 651–668 (2019).

27. P. Monzo, M. Crestani, N. C. Gauthier, “In Vitro Mechanobiology of Glioma: Mimicking the Brain Blood Vessels and White Matter Tracts Invasion Paths” in Brain Tumors (Springer, 2021), pp. 159–196.

28. A. Jain, M. Betancur, G. D. Patel, C. M. Valmikinathan, V. J. Mukhatyar, A. Vakharia, S. B. Pai, B. Brahma, T. J. MacDonald, R. V. Bellamkonda, Guiding intracortical brain tumour cells to an extracortical cytotoxic hydrogel using aligned polymeric nanofibres. Nat Mater 13, 308–316 (2014).

29. S. S. Rao, M. T. Nelson, R. Xue, J. K. DeJesus, M. S. Viapiano, J. J. Lannutti, A. Sarkar, J. O. Winter, Mimicking white matter tract topography using core-shell electrospun nanofibers to examine migration of malignant brain tumors. Biomaterials 34, 5181–5190 (2013).

30. D. Gallego-Perez, N. Higuita-Castro, L. Denning, J. DeJesus, K. Dahl, A. Sarkar, D. J. Hansford, Microfabricated mimics of in vivo structural cues for the study of guided tumor cell migration. Lab Chip 12, 4424–4432 (2012).

31. J. Cha, I. Koh, Y. Choi, J. Lee, C. Choi, P. Kim, Tapered microtract array platform for antimigratory drug screening of human glioblastoma multiforme. Adv Healthc Mater 4, 405–411 (2015).

32. P. Monzo, Y. K. Chong, C. Guetta-Terrier, A. Krishnasamy, S. R. Sathe, E. K. Yim, W. H. Ng, B. T. Ang, C. Tang, B. Ladoux, N. C. Gauthier, M. P. Sheetz, Mechanical confinement triggers glioma linear migration dependent on formin FHOD3. Mol Biol Cell 27, 1246–1261 (2016).

33. J. Cha, S. G. Kang, P. Kim, Strategies of Mesenchymal Invasion of Patient-derived Brain Tumors: Microenvironmental Adaptation. Sci Rep 6, 24912 (2016).

34. T. A. Ulrich, A. Jain, K. Tanner, J. L. MacKay, S. Kumar, Probing cellular mechanobiology in three-dimensional culture with collagen-agarose matrices. Biomaterials 31, 1875–1884 (2010).

35. D. Truong, R. Fiorelli, E. S. Barrientos, E. L. Melendez, N. Sanai, S. Mehta, M. Nikkhah, A three-dimensional (3D) organotypic microfluidic model for glioma stem cells - Vascular interactions. Biomaterials 198, 63–77 (2019).

36. M. T. Ngo, J. N. Sarkaria, B. A. C. Harley, Pericytes and Astrocytes Instruct Glioblastoma Invasion, Proliferation, and Therapeutic Response within an Engineered Brain Perivascular Niche Model. bioRxiv, 2022.2004.2027.489740 (2022).

37. A. Linkous, D. Balamatsias, M. Snuderl, L. Edwards, K. Miyaguchi, T. Milner, B. Reich, L. Cohen-Gould, A. Storaska, Y. Nakayama, E. Schenkein, R. Singhania, S. Cirigliano, T. Magdeldin, Y. Lin, G. Nanjangud, K. Chadalavada, D. Pisapia, C. Liston, H. A. Fine, Modeling Patient-Derived Glioblastoma with Cerebral Organoids. Cell Rep 26, 3203–3211 e3205 (2019).

38. M. A. Marques-Torrejon, E. Gangoso, S. M. Pollard, Modelling glioblastoma tumour-host cell interactions using adult brain organotypic slice co-culture. Dis Model Mech 11, (2018).

39. M. Osswald, E. Jung, F. Sahm, G. Solecki, V. Venkataramani, J. Blaes, S. Weil, H. Horstmann, B. Wiestler, M. Syed, L. Huang, M. Ratliff, K. Karimian Jazi, F. T. Kurz, T. Schmenger, D. Lemke, M. Gommel, M. Pauli, Y. Liao, P. Haring, S. Pusch, V. Herl, C. Steinhauser, D. Krunic, M. Jarahian, H. Miletic, A. S. Berghoff, O. Griesbeck, G. Kalamakis, O. Garaschuk, M. Preusser, S. Weiss, H. Liu, S. Heiland, M. Platten, P. E. Huber, T. Kuner, A. von Deimling, W. Wick, F. Winkler, Brain tumour cells interconnect to a functional and resistant network. Nature 528, 93–98 (2015).

40. P. Monzo, M. Crestani, Y. K. Chong, A. Ghisleni, K. Hennig, Q. Li, N. Kakogiannos, M. Giannotta, C. Richichi, T. Dini, Adaptive mechanoproperties mediated by the formin FMN1 characterize glioblastoma fitness for invasion. Developmental cell, (2021).

41. C. G. Hubert, M. Rivera, L. C. Spangler, Q. Wu, S. C. Mack, B. C. Prager, M. Couce, R. E. McLendon, A. E. Sloan, J. N. Rich, A three-dimensional organoid culture system derived from human glioblastomas recapitulates the hypoxic gradients and cancer stem cell heterogeneity of tumors found in vivo. Cancer research 76, 2465–2477 (2016).

42. F. Polleux, A. Ghosh, The slice overlay assay: a versatile tool to study the influence of extracellular signals on neuronal development. Science’s STKE 2002, pl9–pl9 (2002).

43. M. Vinci, S. Gowan, F. Boxall, L. Patterson, M. Zimmermann, W. Court, C. Lomas, M. Mendiola, D. Hardisson, S. A. Eccles, Advances in establishment and analysis of three-dimensional tumor spheroid-based functional assays for target validation and drug evaluation. BMC Biol 10, 29 (2012).

44. J. E. Ron, P. Monzo, N. C. Gauthier, R. Voituriez, N. S. Gov, One-dimensional cell motility patterns. Physical Review Research 2, 033237 (2020).

45. J. Johnson, M. O. Nowicki, C. H. Lee, E. A. Chiocca, M. S. Viapiano, S. E. Lawler, J. J. Lannutti, Quantitative analysis of complex glioma cell migration on electrospun polycaprolactone using time-lapse microscopy. Tissue Eng Part C Methods 15, 531–540 (2009).

46. C. Beadle, M. C. Assanah, P. Monzo, R. Vallee, S. S. Rosenfeld, P. Canoll, The role of myosin II in glioma invasion of the brain. Mol Biol Cell 19, 3357–3368 (2008).

47. C. Neftel, J. Laffy, M. G. Filbin, T. Hara, M. E. Shore, G. J. Rahme, A. R. Richman, D. Silverbush, M. L. Shaw, C. M. Hebert, An integrative model of cellular states, plasticity, and genetics for glioblastoma. Cell 178, 835–849. e821 (2019).

48. S. Piccirillo, R. Combi, L. Cajola, A. Patrizi, S. Redaelli, A. Bentivegna, S. Baronchelli, G. Maira, B. Pollo, A. Mangiola, Distinct pools of cancer stem-like cells coexist within human glioblastomas and display different tumorigenicity and independent genomic evolution. Oncogene 28, 1807–1811 (2009).

49. J. Klughammer, B. Kiesel, T. Roetzer, N. Fortelny, A. Nemc, K.-H. Nenning, J. Furtner, N. C. Sheffield, P. Datlinger, N. Peter, The DNA methylation landscape of glioblastoma disease progression shows extensive heterogeneity in time and space. Nature medicine 24, 1611–1624 (2018).

50. X. Lu, N. P. Maturi, M. Jarvius, I. Yildirim, Y. Dang, L. Zhao, Y. Xie, E. Tan, P. Xing, R. Larsson, Cell-lineage controlled epigenetic regulation in glioblastoma stem cells determines functionally distinct subgroups and predicts patient survival. Nature Communications 13, 1–16 (2022).

51. S. Darmanis, S. A. Sloan, D. Croote, M. Mignardi, S. Chernikova, P. Samghababi, Y. Zhang, N. Neff, M. Kowarsky, C. Caneda, Single-cell RNA-seq analysis of infiltrating neoplastic cells at the migrating front of human glioblastoma. Cell reports 21, 1399–1410 (2017).

52. J. Wang, E. Cazzato, E. Ladewig, V. Frattini, D. I. Rosenbloom, S. Zairis, F. Abate, Z. Liu, O. Elliott, Y. J. Shin, J. K. Lee, I. H. Lee, W. Y. Park, M. Eoli, A. J. Blumberg, A. Lasorella, D. H. Nam, G. Finocchiaro, A. Iavarone, R. Rabadan, Clonal evolution of glioblastoma under therapy. Nat Genet 48, 768–776 (2016).

53. V. G. LeBlanc, D. L. Trinh, S. Aslanpour, M. Hughes, D. Livingstone, D. Jin, B. Y. Ahn, M. D. Blough, J. G. Cairncross, J. A. Chan, Single-cell landscapes of primary glioblastomas and matched explants and cell lines show variable retention of inter-and intratumor heterogeneity. Cancer Cell 40, 379–392. e379 (2022).

54. M. Fenech, V. Girod, V. Claveria, S. Meance, M. Abkarian, B. Charlot, Microfluidic blood vasculature replicas using backside lithography. Lab Chip 19, 2096–2106 (2019).

55. A. D. Doyle, F. W. Wang, K. Matsumoto, K. M. Yamada, One-dimensional topography underlies three-dimensional fibrillar cell migration. Journal of cell biology 184, 481–490 (2009).

56. P. Gritsenko, W. Leenders, P. Friedl, Recapitulating in vivo-like plasticity of glioma cell invasion along blood vessels and in astrocyte-rich stroma. Histochem Cell Biol 148, 395–406 (2017).

57. E. Serres, F. Debarbieux, F. Stanchi, L. Maggiorella, D. Grall, L. Turchi, F. Burel-Vandenbos, D. Figarella-Branger, T. Virolle, G. Rougon, E. Van Obberghen-Schilling, Fibronectin expression in glioblastomas promotes cell cohesion, collective invasion of basement membrane in vitro and orthotopic tumor growth in mice. Oncogene 33, 3451–3462 (2014).

58. M. L. Suvà, I. Tirosh, The glioma stem cell model in the era of single-cell genomics. Cancer cell 37, 630–636 (2020).

59. J. D. Lathia, J. Gallagher, J. M. Heddleston, J. Wang, C. E. Eyler, J. MacSwords, Q. Wu, A. Vasanji, R. E. McLendon, A. B. Hjelmeland, Integrin alpha 6 regulates glioblastoma stem cells. Cell stem cell 6, 421–432 (2010).

60. M. Crestani, T. Dini, N. C. Gauthier, P. Monzo, Protocol to assess human glioma propagating cell migration on linear micropatterns mimicking brain invasion tracks. STAR protocols 3, 101331 (2022).

61. E. E. Er, M. Valiente, K. Ganesh, Y. Zou, S. Agrawal, J. Hu, B. Griscom, M. Rosenblum, A. Boire, E. Brogi, Pericyte-like spreading by disseminated cancer cells activates YAP and MRTF for metastatic colonization. Nature cell biology 20, 966–978 (2018).

62. Y. Shin, S. Han, J. S. Jeon, K. Yamamoto, I. K. Zervantonakis, R. Sudo, R. D. Kamm, S. Chung, Microfluidic assay for simultaneous culture of multiple cell types on surfaces or within hydrogels. Nature protocols 7, 1247–1259 (2012).

63. R. Gorelik, A. Gautreau, Quantitative and unbiased analysis of directional persistence in cell migration. Nature protocols 9, 1931–1943 (2014).

64. R. Rezakhaniha, A. Agianniotis, J. T. C. Schrauwen, A. Griffa, D. Sage, C. v. Bouten, F. Van De Vosse, M. Unser, N. Stergiopulos, Experimental investigation of collagen waviness and orientation in the arterial adventitia using confocal laser scanning microscopy. Biomechanics and modeling in mechanobiology 11, 461–473 (2012).

65. A. Dobin, C. A. Davis, F. Schlesinger, J. Drenkow, C. Zaleski, S. Jha, P. Batut, M. Chaisson, T. R. Gingeras, STAR: ultrafast universal RNA-seq aligner. Bioinformatics 29, 15–21 (2013).

66. R. Patro, G. Duggal, M. I. Love, R. A. Irizarry, C. Kingsford, Salmon provides fast and bias-aware quantification of transcript expression. Nature methods 14, 417–419 (2017).

67. M. I. Love, W. Huber, S. Anders, Moderated estimation of fold change and dispersion for RNA-seq data with DESeq2. Genome biology 15, 1–21 (2014).

68. A. Sergushichev, An algorithm for fast preranked gene set enrichment analysis using cumulative statistic calculation. BioRxiv 60012, 1–9 (2016).

