## Supplementary material for "SP2G: an imaging and analysis pipeline revealing the inter and intra-patient migratory diversity of glioblastoma"

Michele Crestani *et al.*

**This PDF file includes:**

Figs. S1 to S5
Movies S1 to S6
Data S1 to S2
Appendix

**Other Supplementary Materials for this manuscript include the following:**

Movies S1 to S6
Data S1 to S2
Appendix

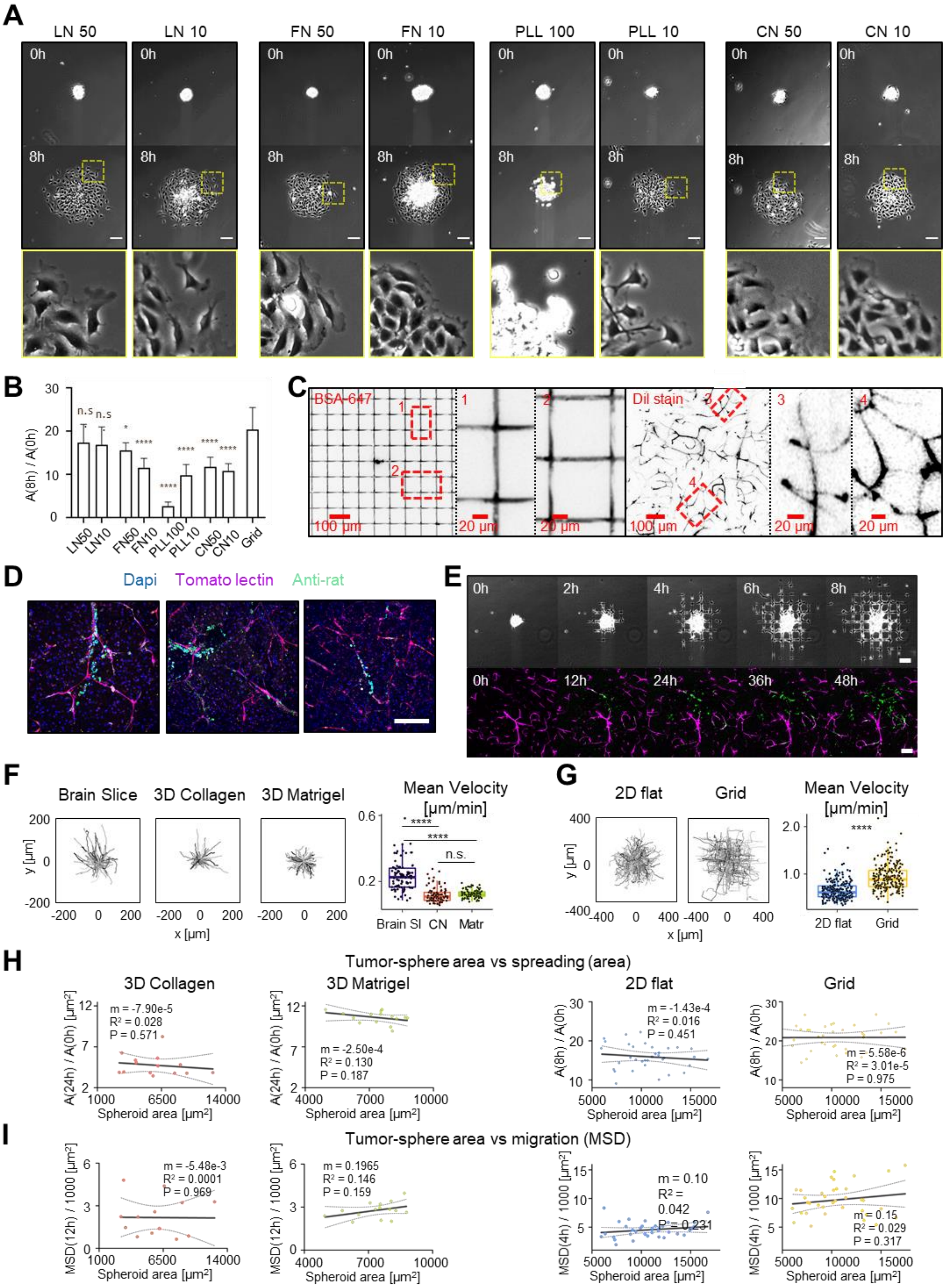
A
LN 50
LN 10
FN 50
FN 10
PLL 100
PLL 10
CN 50
CN 10
0h
8h
B
A(8h) / A(0h)
n.s
n.s
LN50
LN10
FN50
FN10
PLL100
PLL10
CN50
CN10
Grid
C
BSA-647
Dil stain
100 µm
20 µm
D
Dapi
Tomato lectin
Anti-rat
E
0h
2h
4h
6h
8h
12h
24h
36h
48h
F
Brain Slice
3D Collagen
3D Matrigel
Mean Velocity [µm/min]
y [µm]
x [µm]
n.s.
Brain Sl
CN
Matr
G
2D flat
Grid
H
Tumor-sphere area vs spreading (area)
A(24h) / A(0h) [µm²]
A(8h) / A(0h)
Spheroid area [µm²]
m = -7.90e-5
R² = 0.028
P = 0.571
m = -2.50e-4
R² = 0.130
P = 0.187
m = -1.43e-4
R² = 0.016
P = 0.451
m = 5.58e-6
R² = 3.01e-5
P = 0.975
I
Tumor-sphere area vs migration (MSD)
MSD(12h) / 1000 [µm²]
MSD(4h) / 1000 [µm²]
m = -5.48e-3
R² = 0.0001
P = 0.969
m = 0.1965
R² = 0.146
P = 0.159
m = 0.10
R² =
0.042
P = 0.231
m = 0.15
R² = 0.029
P = 0.317

**Fig. S1. SP2G mimics glioblastoma invasion on brain blood vessels.** (**A-B**) Rat C6 glioma cells cultured as spheroids were seeded on glass bottom dishes coated with laminin (LN), fibronectin (FN), Poly-L-Lysine (PLL) and collagen (CN) at 10, 50 or 100 μg/ml and imaged for 8h. (**A**) First and last images of the movies (upper panels) and zoom at 8h (lower panels). (**B**) Quantification of spheroid spreading (area at 8 h over the initial area). Error bars are SD. n = 10, 7, 8, 6, 8, 8, 8, 8, 12 spheroids. One-way ANOVA, p<0.0001. Each condition was compared to the grid condition. **(C)** Comparison of gridded micropatterns (stained with fluorescent BSA, left) with the mouse brain vasculature (stained with Dil dye, right). **(D)** Mouse brain slices invaded by rat C6 glioma cells for 48h, fixed and stained with DAPI, tomato lectin (blood vessels) and anti-rat antibody. **(E)** Montage extracted from time lapse movies of C6 spheroid spreading on gridded micropatterns (top) and on brain slice (bottom). **(F)** Cell trajectories and mean velocities of C6 single cells migrating in brain slice, 3D collagen and 3D Matrigel. n = 80, 95, 90 tracks; 5 to 7 tracks per spheroid, each dot is a cell. One-way ANOVA. **(G)** Cell trajectories and mean velocities of single cells migrating on 2D flat and gridded micropatterns. n = 215, 215 tracks; 5 to 7 tracks per spheroid, each dot is a cell. Unpaired t-test. **(H-I)** Spheroid spreading (Afinal/A0) and Mean Squared Displacement (MSD) measured in function of the spheroid area at the initial time. Line slope (m), coefficient of determination ($R^2$) and p-value (P) of the linear regression are reported. Pearson's r correlation coefficients, from left to right: -0.16, -0.36, -0.13, 0.01. (n = 14, 15, 35, 35 spheroids)(H) and -0.01, 0.38, 0.17, 0.20 (n = 14, 15, 35, 35 spheroids) (I).

### A. Migration area, A(t). *Semi-automated step.*

Place raw data in a folder. Raw data can be:
- Several bio-format files (e.g. 1 per condition)
- .tif files (1 per spheroid) placed in a sub-folder

Run code "*Main: Polygon Tracking and Segmentation*". Inputs: raw data. Outputs are:

1 folder per raw data file, in which 1 subfolder per sample contains:
- Binarized grid
- Binarized cells
- Polygonal shape
- Binarized grid nodes
- .avi movie
- .txt file (debugging)

- Run code "*Average Polygon*". Input: *polygonal shape*.
- Output: graphical representation for A(t)

### B. Diffusivity, D = δA(t) / δt; Boundary speed, $v(t) = D(t) / (2\sqrt{(A(t))})$. *Automated step.*

Run code "*Polygon Area Measurement*". Input: *polygonal shape*. Outputs, in the same .csv file :
- Migration areas A(t) (absolute and relative to $t_0$)
- Smoothed A(t)
- Diffusivity D(t)
- Boundary speed v(t)

### C. Extrapolate τ and Δ from mean A(t) and $v(t)$. *Semi-automated/User step.*

- τ normalizes over migration area. It is the time step when the mean A(t) reaches the lowest A(final_time) among the experimental conditions to compare.
- Δ normalizes over boundary speed. It is the time window taken by the cells to travel 100 μm. If Δ < 8h, cells are considered motile. Δ must be lower than τ.

*See supplementary file 1, it can be utilized as template.*

### D. Collective migration and Directional Persistence. *Automated step.*

Run code "*Collective migration and Directional Persistence*". Input: *binarized cells*. Output:

Collective migration and Directional Persistence values in .csv file (mean and indexed by spheroid name. All values indexed by time point.)

### E. Hurdling. *Semi-automated step.*

Run code "*Hurdling*". Input: *binarized cells* and *binarized grid*. Outputs:
- Average Intensities (1 value per square) + Mean Average Intensity (1 value) in .csv file, for relative hurdling calculation.
- Average intensities (1 value per square) normalized over the maximum value of the dataset in the same .csv file, for visualizing the cumulative distribution.

All values indexed by time point. *See supplementary file 2.*

F G

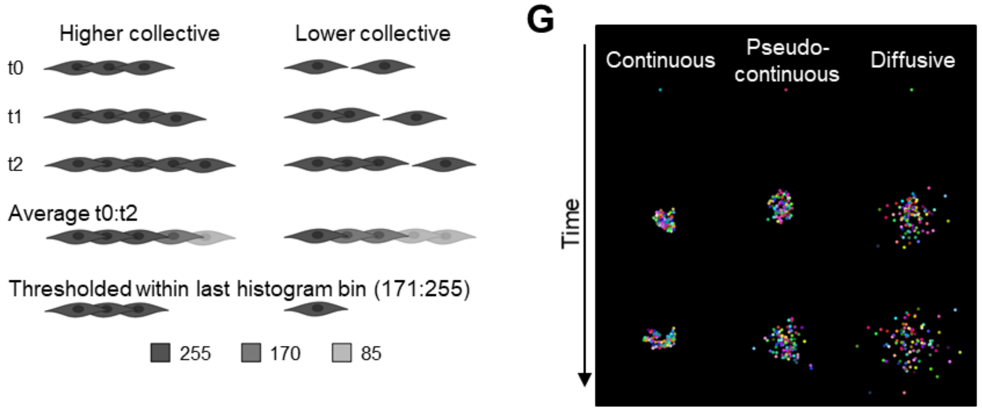


H

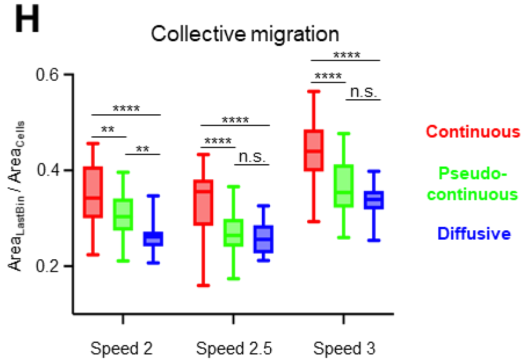


I

| Speed | Diffusion | τ−Δ | Δ |
|---|---|---|---|
| 2 | *Continuous* | 285 | 15 |
| | *Pseudo-cont* | 105 | |
| | *Diffusive* | 49 | |
| 2.5 | *Continuous* | 93 | 12 |
| | *Pseudo-cont* | 69 | |
| | *Diffusive* | 31 | |
| 3 | *Continuous* | 69 | 10 |
| | *Pseudo-cont* | 49 | |
| | *Diffusive* | 20 | |

**Fig. S2. SP2G: from images to quantitative data. (A-E)** Table summarizing SP2G quantitative outputs. **(F)** Schematic representation of the rationale allowing collective migration quantification. Average images obtained from highly collective cells (left) will return higher values and form higher area when thresholded, compared to low collective cells (right). **(G-I)** SP2G outputs obtained with simulated data using 100 round particles (radius=5 pixels) diffusing with full constraint (Continuous, 100% probability of being attached to a neighbor), partial constraint (Pseudo-continuous, 90% probability of being attached to a neighbor), or no constraint (Diffusive, simple diffusion), over 300 time frames at 3 speed regimes, with mean speed of 2, 2.5, 3 pixels per frame and standard deviation of 2, 2.5, 3 pixels, respectively (9 conditions total). **(G)** Example of output obtained with speed = 3±3 pixels per frame. **(H)** Collective migration output obtained in the 9 conditions. **(I)** Table highlighting each simulated condition, their corresponding $\Delta$ ($\Delta$ was chosen so as its product with speed equals 30) and $\tau$ - $\Delta$. One-way ANOVA, n=30 simulations per condition.

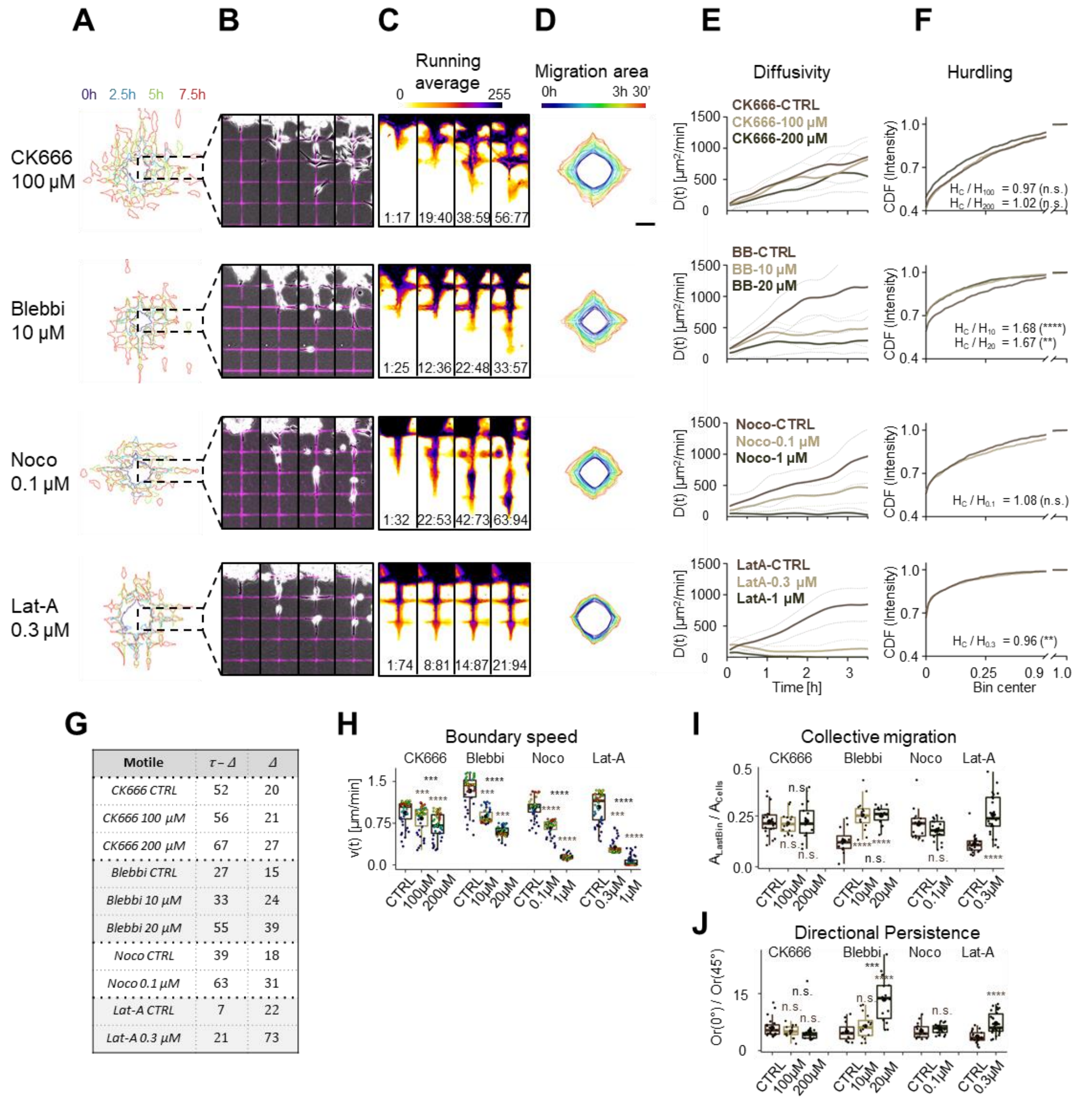


| Motile | τ − Δ | Δ |
|---|---|---|
| CK666 CTRL | 52 | 20 |
| CK666 100 μM | 56 | 21 |
| CK666 200 μM | 67 | 27 |
| Blebbi CTRL | 27 | 15 |
| Blebbi 10 μM | 33 | 24 |
| Blebbi 20 μM | 55 | 39 |
| Noco CTRL | 39 | 18 |
| Noco 0.1 μM | 63 | 31 |
| Lat-A CTRL | 7 | 22 |
| Lat-A 0.3 μM | 21 | 73 |

**Fig. 3. SP2G quantifies cell migratory tactic changes upon cytoskeleton drugs.** Patient-derived glioblastoma spheroids, NNI-21, were seeded on fluorescent gridded micropatterns, imaged for 8h in presence of the drugs, and analyzed as indicated in fig 2 (number of spheroids: n= 16, 14, 16, 14, 20, 12, 26, 16 for CK666 100 and 200 μM, blebbistatin 10 and 20 μM, Nocodazole 0.1 and 1 μM, latrunculin-A 0.3 and 1 μM respectively and n = 25, 12, 16, 21 for their respective control, n = 2 independent experiments per condition. **(A-C)** Cellular edges (A) and corresponding overlays of the phase contrast and the fluorescent grid images (B) at 4 time points (0h, 2.5h, 5h, 7.5h) and corresponding running average (RA) (C). The time window Δ constituting the corresponding RA frame is indicated at the bottom of each panel. **(D)** Average polygon visualizing migration area. **(E)** Diffusivity over 3 h 30' (from top to bottom: p=0.0003, p<0.0001, p<0.0001, p<0.0001 Kruskal-Wallis test). Dashed lines are standard deviation. Dunn's multiple comparison test: CK666: CTRL vs 100 and 100 vs 200 n.s, CTRL vs 200 p=0.0002; others: p<0.0001 for all, except

CTRL vs lat-A 0.1 μM p=0.0007. (**F**) Hurdling visualized as the Cumulative Distribution Function (CDF) of the normalized mean intensity of the grid squares. Kolmogorov-Smirnov tests: CK666-CTRL vs CK666-100 p=0.9926 (n.s.), CK666-CTRL vs CK666-200 p=0.6828 (n.s.), BB-CTRL vs BB10 p=5.3505e-05 (****), BB-CTRL vs BB-20 p=0.0019 (**) , Noco-CTRL vs Noco-0.1 p=0.3499 (n.s.), LatA-CTRL vs LatA-0.3 p=0.0087 (**). (**G**) Δ (number of frames needed to travel 100 μm) and τ - Δ (number of frames in the RA movie) in motile cells (faster than 100 μm / 8 h = 0.21 μm/min) (**H**) Mean boundary speed over 3h 30'. Each dot represents a time-point and is color-coded as in D (p<0.0001, Kruskal-Wallis test with Dunn's multiple comparison test. (**I**) Collective migration for the motile cells. Each dot represents a spheroid. One-way ANOVA for CK666 (n.s.) and blebbistatin (****) with Dunn's multiple comparison test. t-test for nocodazole and latrunculin-A. (**J**) Directional persistence for the motile cells. Each dot represents a spheroid. One-way ANOVA for CK666 (n.s.) and blebbistatin (****) with Dunn's multiple comparison test. t-test for nocodazole and latrunculin-A. Scale bars are 100 μm. In all the boxplots, the middle horizontal line represents the median and the black dot is the mean value. The stars above each treated condition compare it against the control, the stars in middle-top compare the 2 treated conditions. *** means p<0.001, **** means p<0.0001. Time and image intensity are color-coded as indicated.

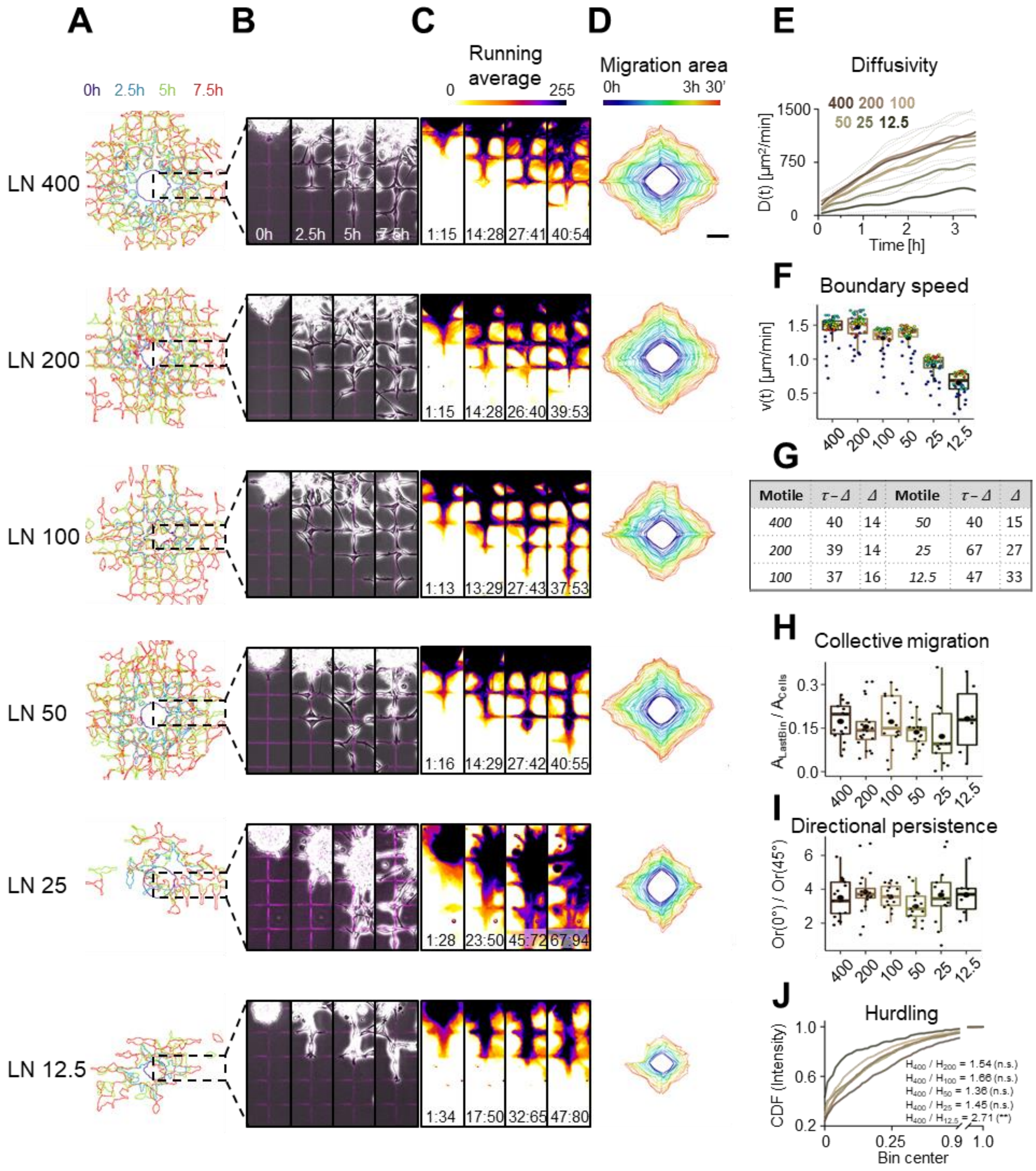


**Fig. S4. SP2G detects cell migratory tactic changes upon variations of laminin concentration.** Analysis of NNI-21 spheroid spreading on gridded micropatterns printed with laminin (LN) at 400, 200, 100, 50, 25, 12.5 μg/ml (n = 17, 20, 16, 17, 13, 6 spheroids, n = 3 independent experiments per condition). **(A-C)** Cellular edges (A) and corresponding overlays of the phase contrast and the fluorescent grid images (B) at 4 time points and corresponding running average (RA) (C). **(D)** Average polygon visualizing migration area. **(E)** Diffusivity over 3 h 30’ ($p<0.0001$, Kruskal-Wallis test). Dashed lines are the standard deviation. Dunn’s multiple comparison test:

400 vs 25 / 12.5, 200 vs 25 / 12.5 and 100 / 50 vs 12.5 p<0.0001; 100 / 50 vs 25 and 25 vs 12.5 p<0.05; others n.s. **(F)** Mean boundary speed over 3h 30’. Each dot represents a time-point and is color-coded as in D (p<0.0001, Kruskal-Wallis test for all). Dunn’s multiple comparison test: 400 vs 25 / 12.5, 200 vs 25 / 12.5 and 100 / 50 vs 12.5 p<0.0001; 100 vs 25 p=0.0001; 200 vs 100 p=0.0086; 400 vs 100 / 50 and 200 vs 50 p<0.05; others n.s. **(G)** $\Delta$ and $\tau$ - $\Delta$ in motile cells (faster than 0.21 μm/min). **(H)** Collective migration for the motile cells. Each dot represents a spheroid. One-way ANOVA (n.s.) with Dunn’s multiple comparison test (all n.s.). **(I)** Directional persistence for the motile cells. Each dot represents a spheroid. One-way ANOVA (n.s.) with Dunn’s multiple comparison test (all n.s.). **(J)** Hurdling visualized as the Cumulative Distribution Function (CDF) of the normalized mean intensity of the grid squares. Kolmogorov-Smirnov tests: LN400 vs LN200 p=0.4496 (n.s.), LN400 vs LN100 p=0.1976 (n.s.), LN400 vs LN50 p=0.4496 (n.s.), LN400 vs LN25 p=0.2659 (n.s.), LN400 vs LN12.5 p=0.0054 (**). Scale bars are 100 μm. In all the boxplots, the middle horizontal line represents the median and the black dot is the mean value. Time and image intensity are color-coded as indicated.

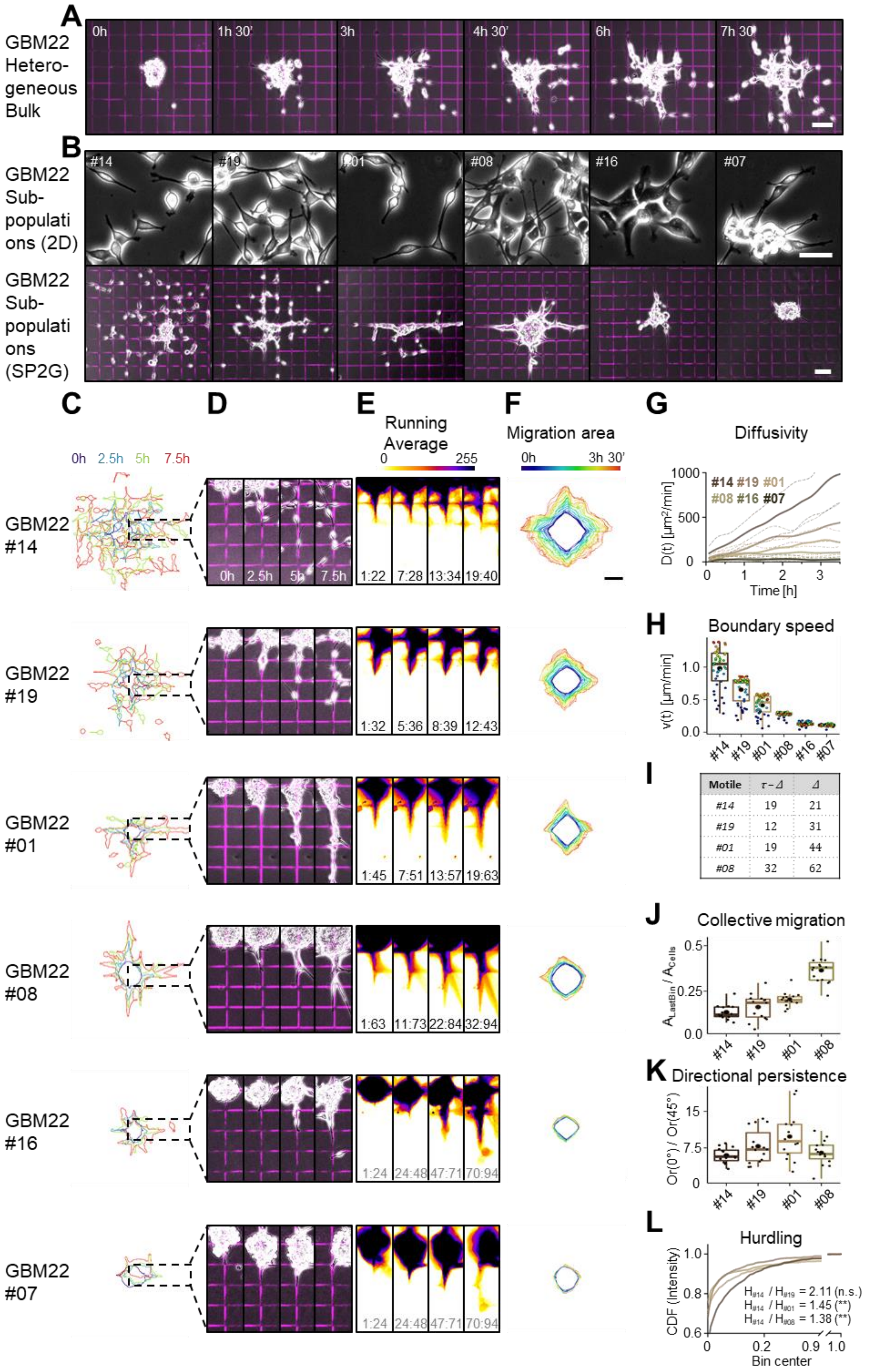
A
GBM22 Hetero-geneous Bulk
0h
1h 30'
3h
4h 30'
6h
7h 30'
B
GBM22 Sub-populati ons (2D)
#14
#19
#01
#08
#16
#07
GBM22 Sub-populati ons (SP2G)
C
D
E
F
G
0h 2.5h 5h 7.5h
Running Average
0 255
Migration area
0h 3h 30'
Diffusivity
GBM22 #14
GBM22 #19
GBM22 #01
GBM22 #08
GBM22 #16
GBM22 #07
1:22 7:28 13:34 19:40
1:32 5:36 8:39 12:43
1:45 7:51 13:57 19:63
1:63 11:73 22:84 32:94
1:24 24:48 47:71 70:94
#14 #19 #01 #08 #16 #07
D(t) [μm²/min]
Time [h]
H
Boundary speed
v(t) [μm/min]
I
Motile τ−Δ Δ
#14 19 21
#19 12 31
#01 19 44
#08 32 62
J
Collective migration
A_LastBin / A_Cells
K
Directional persistence
Or(0°) / Or(45°)
L
Hurdling
CDF (Intensity)
Bin center
H#14 / H#19 = 2.11 (n.s.)
H#14 / H#01 = 1.45 (**)
H#14 / H#08 = 1.38 (**)

**Fig. S5. SP2G reveals heterogeneity in the migratory tactics adopted by glioblastoma sub-populations isolated from patient-derived cell lines.** Spheroids from GBM22 original cell line and isolated subpopulations (clones #14, #19, #01, #08, #16, #07) were seeded on fluorescent gridded micropatterns, imaged for 8h, and analyzed as indicated in fig 2. **(A)** Snapshots extracted from the movie of the original population at the indicated time-points. **(B)** Phase-contrast pictures of the GBM7 sub-populations cultured on laminin (top panel, bar is 50 μm) and after SP2G at 8 hours (bottom, bar is 100 μm). **(C-E)** SP2G analysis of the clones #14, #19, #01, #08, #16, #07 (n = 15, 15, 14, 15, 13, 11 spheroids. n = 2 independent experiments): Cellular edges (C), overlays of the phase contrast and the fluorescent grid images at 4 time points (D) and corresponding running average (RA) (E). **(F)** Average polygon visualizing migration area. **(G)** Diffusivity over 3 h 30' ($p<0.0001$, Kruskal-Wallis test). Dashed lines are the standard deviation. Dunn's multiple comparison test: #14 vs #01 / #08 / #16 / #07, #19 vs #16 / #07, #01 vs #16 / #07, #08 vs #16 / #07 $p<0.0001$; #19 vs #08 $p=0.0051$; others n.s. **(H)** Mean boundary speed over 3h 30'. Each dot represents a time-point and is color-coded as in F ($p<0.0001$, Kruskal-Wallis test). Dunn's multiple comparison test: #14 vs #01 / #08 / #16 / #07, #19 vs #16 / #07, #01 vs #16 / #07, #08 vs #07 $p<0.0001$; #19 vs #08 $p=0.0001$; #08 vs #16 $p=0.0017$; others n.s. **(I)** $\Delta$ and $\tau$ - $\Delta$ of the motile subpopulations. **(J)** Collective migration for the motile cells #14, #19, #01 and #08. Each dot represents a spheroid. $p<0.0001$, ordinary one-way ANOVA. Multiple comparisons: #08 vs #14 / #19 / #01 $p<0.0001$; #01 vs #14 $p=0.0085$; others n.s. **(K)** Directional persistence for the motile #14, #19, #01 and #08. Each dot represents a spheroid. $p=0.0134$, ordinary one-way ANOVA. Multiple comparisons: #01 vs #08 / #14 $p<0.05$; others n.s. **(L)** Hurdling visualized as the Cumulative Distribution Function (CDF) of the normalized mean intensity of the grid squares. Kolmogorov-Smirnov tests: #14 vs #19 $p=0.0715$ (n.s.), #14 vs #01 $p=0.0019$ (**), #14 vs #08 $p=0.0011$. The ratio indicates the relationship between the average mean intensities of the most hurdling (#14) against the others. Bars are 100 μm (A, B bottom panel, F) and 50 μm (B, top panel). In all the boxplots, the middle horizontal line represents the median and the black dot is the mean value. Time and image intensity are color-coded as indicated.

**Movie S1.**

Concatenated timelapse movies of spreading C6 spheroids. 1) brain slices; 2) 3D collagen and matrigel; 3) 2D flat and gridded micropatterns; 4) representative single cells moving in each setup.

**Movie S2.**

Tutorial for SP2G macro suite

**Movie S3.**

SP2G known migration phenotypes. 1) SP2G of NNI-11, NNI-21, NNI-24; 2) SP2G tracking; 3) SP2G averaging; 3) SP2G projected averaging; 4) running average for the motile NNI-21 and NNI-24.

**Movie S4.**

SP2G drug treatments (NNI-21). Top: SP2G tracking, bottom: SP2G averaging. 1) CK666; 2) Blebbistatin; 3) Nocodazole; 4) Latrunculin-A.

**Movie S5.**

SP2G in the GBM7 patient derived cell line. 1) Heterogeneous bulk; 2) intra-patient heterogeneity of the 5 clones; 3) SP2G tracking; 4) SP2G averaging; 5) SP2G projected averaging; 6) running average for the motile #09, #01, #07; 7) spreading spheroids in brain slices of the 5 clones.

**Movie S6.**

SP2G in the GBM22 patient derived cell line. 1) Heterogeneous bulk; 2) intra-patient heterogeneity of the 6 clones; 3) SP2G tracking; 4) SP2G averaging; 5) SP2G projected averaging; 6) running average for the motile clones #14, #19, #01, #08.

**Data S1. (separate file)**

Calculations of Tau, Delta, and mean boundary speed starting from the time trend of the relative area occupied by cells. Imposed parameters: motility threshold (does the invasive front travel faster than a given space - in our case 100 microns - in 8 hours? yes/no), frame rate, and total number of frames.

**Data S2. (separate file)**

Calculations of Average Hurdling and Relative hurdling starting from the raw distribution of intensities in the passivated areas of the grid.

**Annex. (separate file)**

Step by step manual for running SP2G macro suite.
